# A network model of the barrel cortex combined with a differentiator detector reproduces features of the behavioral response to single-neuron stimulation

**DOI:** 10.1101/2020.03.30.016261

**Authors:** Davide Bernardi, Guy Doron, Michael Brecht, Benjamin Lindner

**Affiliations:** Bernstein Center for Computational Neuroscience Berlin, Philippstr. 13, Haus 2, 10115 Berlin, Germany; Physics Department of Humboldt University Berlin, Newtonstr. 15, 12489 Berlin, Germany

## Abstract

The stimulation of a single neuron in the rat somatosensory cortex can elicit a behavioral response. The probability of a behavioral response does not depend appreciably on the duration or intensity of a constant stimulation, whereas the response probability increases significantly upon injection of an irregular current. Biological mechanisms that can potentially suppress a constant input signal are present in the dynamics of both neurons and synapses and seem ideal candidates to explain these experimental findings. Here, we study a large network of integrate-and-fire neurons with several salient features of neuronal populations in the rat barrel cortex. The model includes cellular spike-frequency adaptation, experimentally constrained numbers and types of chemical synapses endowed with short-term plasticity, and gap junctions. Numerical simulations of this model indicate that cellular and synaptic adaptation mechanisms alone may not be sufficient to account for the experimental results if the local network activity is read out by an integrator. However, a differentiator circuit can detect the single-cell stimulation with a reliability that barely depends on the length or intensity of the stimulus, but that increases when an irregular signal is used. This finding is in accordance with the experimental results obtained for the stimulation of a regularly-spiking excitatory cell.

**Author summary:** It is widely assumed that only a large group of neurons can encode a stimulus or control behavior. This tenet of neuroscience has been challenged by experiments in which stimulating a single cortical neuron has had a measurable effect on an animal’s behavior. Recently, theoretical studies have explored how a single-neuron stimulation could be detected in a large recurrent network. However, these studies missed essential biological mechanisms of cortical networks and are unable to explain more recent experiments in the barrel cortex. Here, to describe the stimulated brain area, we propose and study a network model endowed with many important biological features of the barrel cortex. Importantly, we also investigate different readout mechanisms, i.e. ways in which the stimulation effects can propagate to other brain areas. We show that a readout network which tracks rapid variations in the local network activity is in agreement with the experiments. Our model demonstrates a possible mechanism for how the stimulation of a single neuron translates into a signal at the population level, which is taken as a proxy of the animal’s response. Our results illustrate the power of spiking neural networks to properly describe the effects of a single neuron’s activity.

## Introduction

A classical method used in neuroscience to understand cortical circuits is to determine how single neurons respond to a controlled sensory stimulus. If, for instance, one of a rat’s whiskers is moved by the experimenter, a change in firing of specific neurons can be measured in the barrel cortex, one of the most well studied parts of the primary sensory cortex [1]. The concept of “reverse physiology” turns this approach around by studying the inverse situation in which neurons in higher brain areas are stimulated and a behavioral or motor response can be elicited (see [2] for early references on such experiments). The two kinds of experiments in combination then allow the linking of sensation and perception, one of the notoriously difficult problems in neuroscience.

In the case that the stimulation affects only a single neuron, the outcome of the unconventional and technically challenging reverse physiology experiments are particularly striking: stimulating a single neuron in the motor cortex can evoke a whisker movement [3], and single-cell stimulation in the barrel cortex (but not in the thalamus) leads to a weak but statistically significant behavioral response [4–6]. This contradicts prevailing hypotheses that relevant signals can only be encoded in the activity of large neural populations.

Both the enormity of cortical networks - tens of thousands of neurons in the case of the somatosensory cortex [7] - and the apparent randomness of single-neuron spiking [8, 9] have classically been evoked as arguments for population coding. If single spikes are unpredictable and noisy how can a few externally induced spikes lead to changes in behavior?

On the theoretical side, cortical populations have been modeled as (locally) random networks of synaptically coupled excitatory and inhibitory cells [10–12] (many studies just take into account two distinct cell types). Even without the inclusion of explicit noise sources, these models can show asynchronous irregular activity [13–15] that is similar to that observed in the cortex of alert animals [16, 17]; this kind of network noise can also be described analytically by stochastic mean-field methods (see for instance [10, 13, 18–20]). Besides the autonomous activity of such networks, their linear and nonlinear response to global stimuli (applied to *all* neurons in the network) has been in focus [20–24]. However, injecting a current into a *single* neuron in such a generic network model can lead to sizable changes in a subpopulation’s activity as well [25, 26]; this subpopulation can be regarded as a readout of the stimulus. If the readout population is somewhat oriented towards the direct postsynaptic partners of the stimulated cell, then the stimulus can be detected in the activity of this population [25]. If, more realistically, the readout is accomplished by a second recurrent network with feed-forward inhibition (as is most likely the case in the cortex), already a very small bias will lead to a detection performance comparable to that in the behavioral experiment [26]. Especially challenging for theoreticians are the results of the nanostimulation experiments in the barrel cortex of behaving rats [6] which demonstrated striking dependencies of the behavioral response on the properties of the stimulating current. The response does not depend on the duration of the stimulus, it depends weakly on its intensity, and strongly on its irregularity: the response is greatly enhanced if the current varies irregularly within the stimulation window instead of being held constant.

Here we tackle these challenges by studying a more detailed computational model that is much closer to the barrel cortex network than the generic setups previously investigated in the context of single-cell stimulation [25, 26]. We also put forward a novel implementation of an alternative readout mechanism, based on the differentiation of the population activity. Our model includes three distinct populations of cells (one excitatory neuron type and two distinct inhibitory interneuron types) that are all modeled as integrate-and-fire neurons endowed with adaptation currents, mediating the omnipresent spike-frequency adaptation. These cells are coupled by chemical synapses, which undergo synaptic depression or facilitation, and by electrical synapses (gap junctions). Combined with the so-called *differentiator readout*, we achieve a response at the population level that is in several key aspects similar to the behavioral response in the experiments by [6].

## Results

The model consists of two parts: a recurrent network, in which a randomly selected excitatory regular spiking cell is stimulated to mimic the experiments, and a readout, which receives input from excitatory neurons of the recurrent network and can detect the stimulation.

### Recurrent network model

Figure 1 shows a scheme containing all essential features of the BCN, briefly described in the figure caption. The recurrent network model represents the surroundings of the stimulated cell in a radius of about 200 μm, in which connection probabilities can be considered as constant [27, 28]. A total network size of *N* = 2600 is in line with estimates of neuron density in the barrel cortex [7]. Although this network size corresponds to a fraction of one barrel, this part of the model will be indicated as “barrel cortex network” (BCN). The BCN consists of three neuronal populations: excitatory regular spiking cells (RS), inhibitory fast-spiking (FS) cells, and somatostatin-expressing low-threshold spiking (SOM-LTS) inhibitory neurons. These three cell types account for a large fraction of the neurons found in the barrel cortex (about 99% of layer IV [29]). All neurons are modeled as one-compartment leaky integrate-and-fire (LIF) neurons. Besides leak conductance and spike generation, several other biological mechanisms are modeled, according to the cell type.

**Fig 1.**
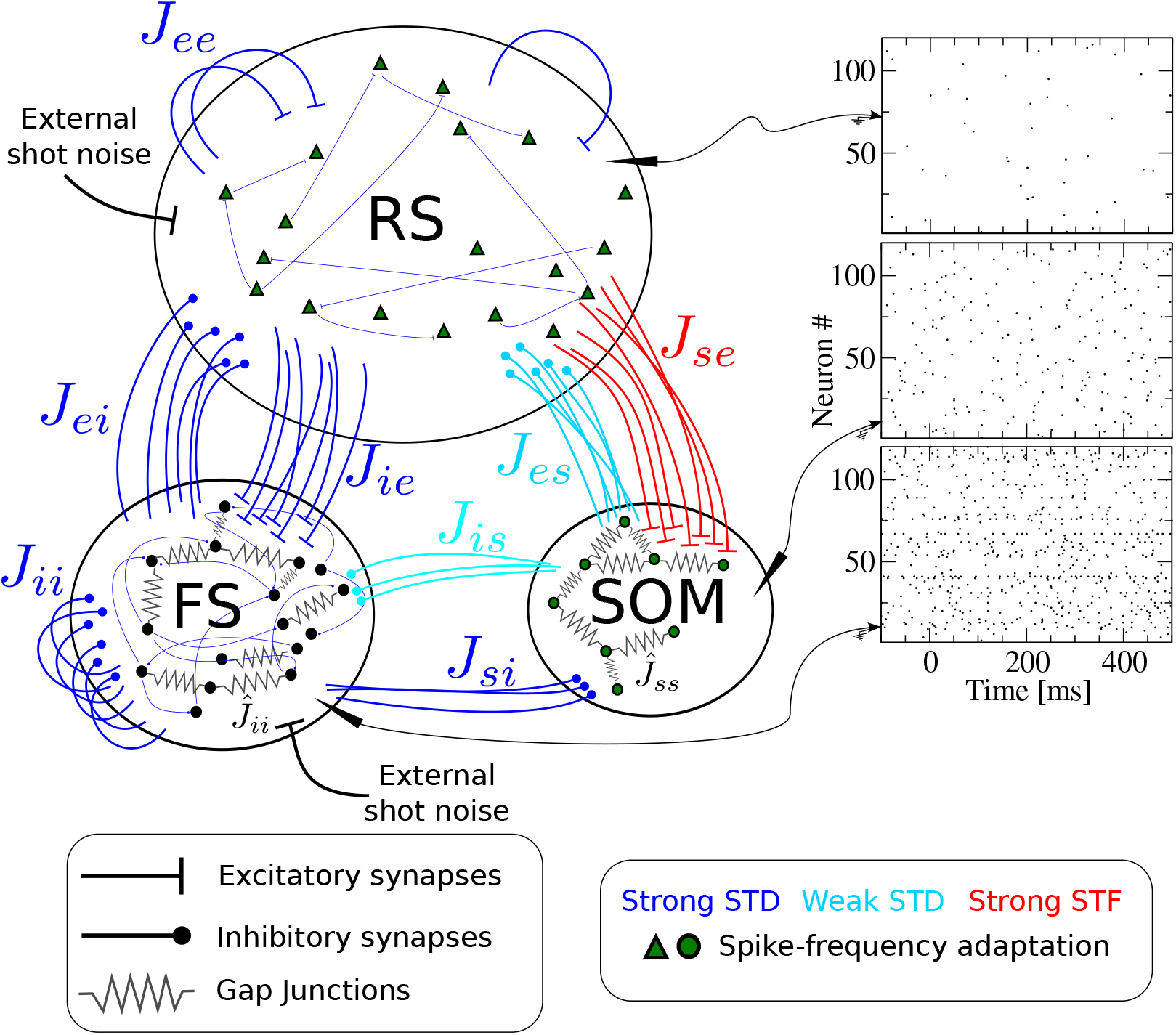
Recurrent network model representing the surroundings of the stimulated cell. The network is formed by *N_e_* = 2000 excitatory regular spiking (RS) neurons, *N_i_* = 400 inhibitory fast spiking (FS) neurons, and *N_s_* = 200 inhibitory somatostatin-positive low-threshold spiking (SOM-LTS) neurons. Recurrent connections between RS neurons are sparse (15%), all connections involving FS neurons as well as those between RS and SOM-LTS neurons are dense (40%-50%). FS and SOM-LTS neurons are electrically coupled (only neurons of the same type). Gap junctions are represented by an effective all-to-all spiking coupling (see main text). Connections in blue are strongly depressing, connections in light blue are weakly depressing, and connections in red are strongly facilitating. RS and SOM-LTS neurons are endowed with a spike-frequency adaptation current. Input from the thalamus and from neighboring cortical regions is represented by Poissonian shot noise. SOM-LTS neurons do not receive external shot noise. The three raster plots show the spontaneous activity of 120 (from top to bottom) RS, SOM, and FS neurons. The spontaneous activity of all three populations is asynchronous and irregular. The spontaneous mean firing rate of excitatory RS neurons is *r*_sp,e_ ≈ 0.8 Hz, of SOM-LTS neurons is *r*_sp,s_ ≈ 3 Hz, and of FS neurons is *r*_sp,i_ ≈ 10 Hz.

The largest population consists of 2000 excitatory RS neurons, which are sparsely connected to each other but densely connected to FS interneurons and SOM-LTS cells, as experimental studies report [29]. The membrane time constant of RS cells is lognormally distributed with mean *τ*_*m,e*_ = 20 ms and a standard deviation of 20% of the mean [29, 30]. The 400 inhibitory FS interneurons are characterized by faster membrane time constants (lognormal distribution with mean *τ*_*m,i*_ = 10 ms [29]) and are densely connected both to other FS neurons and to RS cells. The 200 SOM-LTS inhibitory neurons possess longer time scales (lognormal distribution with mean *τ*_*m,s*_ = 20 ms [29]) and a firing threshold which is 6 mV lower than in RS and FS neurons [29]. SOM-LTS neurons do not inhibit each other via chemical synapses, but form dense connections to and from RS neurons and sparser connections to and from FS interneurons [29, 31, 32].

Both FS and SOM-LTS neurons are densely coupled to cells of the same type via electrical synapses (gap junctions) [29, 31, 33], which are represented here by an effective global excitatory spiking coupling (the sub-threshold contribution of gap junctions has a much smaller effects on the network dynamics [34]).

In the barrel cortex, both RS and SOM-LTS neurons display spike-frequency adaptation, whereas FS neurons do not [29, 35]. Therefore, RS and SOM-LTS are endowed with a spike-triggered hyperpolarizing current [36–38]. Consistent with experimental observations, the strength of the adaptation current is larger for RS neurons than for SOM-LTS neurons.

We drive RS and FS cells with Poissonian spike trains mimicking input from the thalamus and neighboring cortical areas. SOM-LTS cells are mostly subject to local input [29, 39] and therefore, in our model, they do not receive external input.

Experimental studies suggest that most synapses in the barrel cortex show short term depression, with the notable exception of connections from RS neurons to SOM-LTS cells, which have been found to be strongly facilitating [29, 40–42]. The short term plasticity of chemical synapses is simulated here by means of a standard model [43, 44]. Parameters were chosen such that all synapses except those from and to SOM-LTS neurons are strongly depressing. These synapses are depicted in blue in fig. 1. Parameters for synapses branching from SOM-LTS neurons (represented in light blue in fig. 1) are set such that they display a weak depression. Finally, parameters of synapses connecting RS cells to SOM-LTS neurons are chosen to generate a strong facilitation. A further property that distinguishes these synapses is that the transmission failure rate is high (~ 50%) at low presynaptic firing rate, but the reliability increases upon repeated activation of the synapse [29]. This property is modeled by a variable that mimics the activity-dependent failure rate.

More details on model equations and parameters are given in the Methods.

### Readout models

Because outgoing long-range connections in the cortex mostly originate from pyramidal neurons, we assume that the readout can access a subset of the RS cells and consider three possible schemes, as illustrated in fig. 2. The first detection scheme (fig. 2A) receives input from a subset of the excitatory neurons of the BCN and reacts when the filtered activity of these neurons reaches a lower barrier. This readout scheme will be called integrator readout (IR). The second readout scheme (fig. 2B) filters the activity in the same way as the IR, but it subtracts a time-shifted copy of the same activity. In other words, it considers the difference between the filtered activity at different time points, thus acting as a sort of differentiator. For this reason, it will be referred to as differentiator readout (DR). The third readout scheme is the implementation of the DR by means of a simple network of LIF neurons and will be called differentiator network readout (DNR).

**Fig 2.**
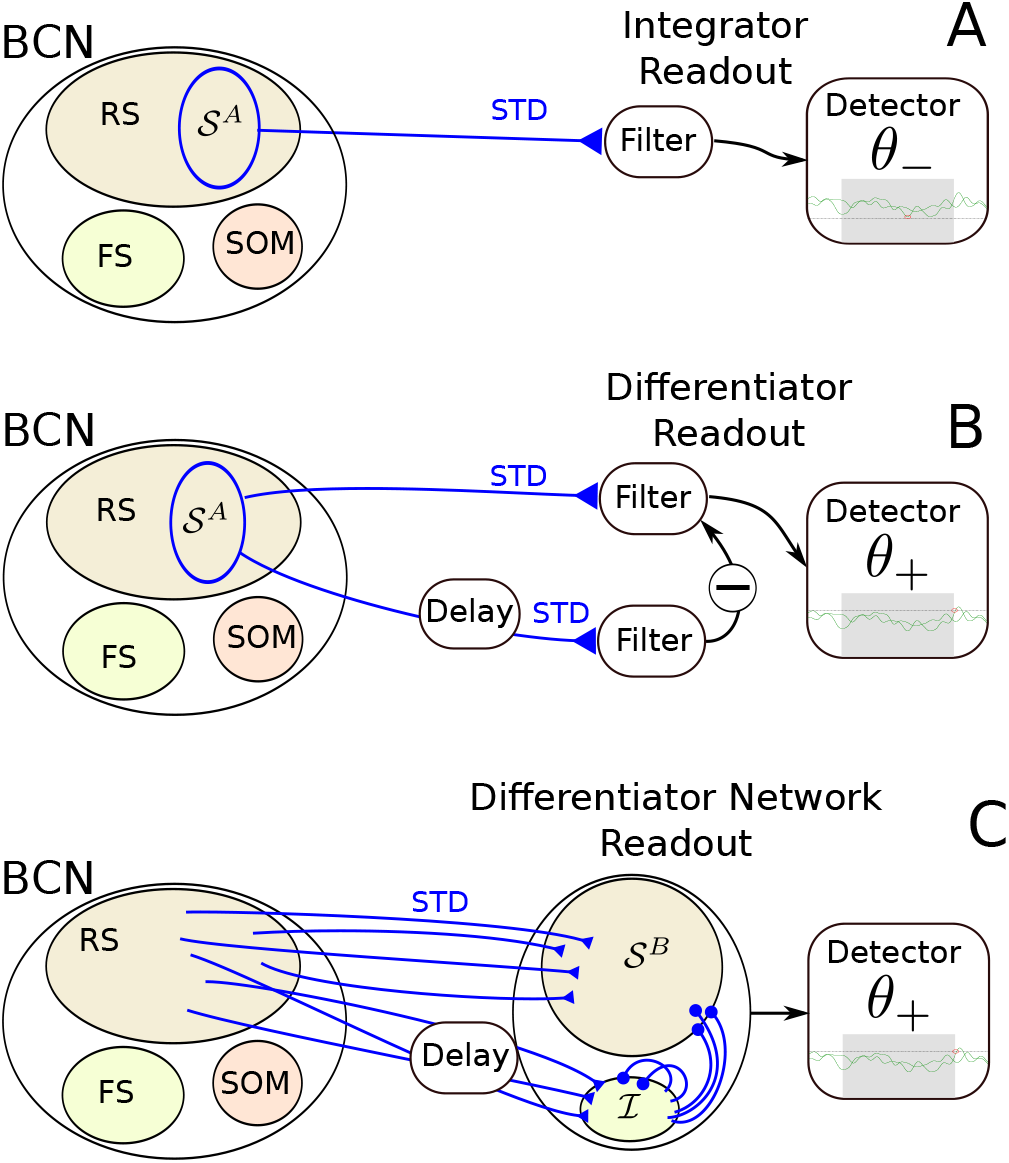
Readout models considered in this paper. A cell selected at random from the barrel cortex network (BCN) is selected at random and stimulated. The BCN consists of three populations: excitatory regular-spiking neurons (RS), inhibitory fast-spiking neurons (FS), and somatostatin-positive low-threshold spiking neurons (SOM-LTS), all modeled by leaky integrate-and-fire neurons. The BCN includes several biological details of the barrel cortex (see fig. 1 and methods). Three readout schemes are considered. **A**: the integrator readout (IR) integrates the activity of a subset of the RS neurons within the BC and reacts to deviations in the negative direction. **B**: the differentiator readout (DR) evaluates the difference between the IR activity at two time points separated by a delay. This filtered running difference at fixed lag is processed by the detector, which reacts when an upper threshold is reached. **C**: the differentiator network readout (DNR) implements the operation of the DR with two populations of LIF neurons. The FS readout population provides delayed recurrent inhibition to itself and feed-forward inhibition to the RS readout population. All connections depicted in blue are dynamic and show short-term depression (STD).

#### Integrator readout

The first readout scheme, the integrator readout (IR), is based on a random selection of 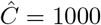 neurons that constitute the readout set 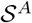 (see fig. 2A). The spike trains emitted by all neurons within the readout population 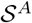 are linearly filtered by using the following dynamical equation:

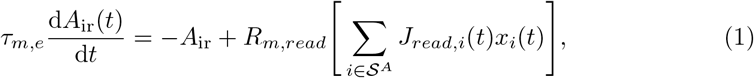

where *x*_*i*_ is the spike train of the *i*th neuron within the readout set 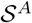, the integration time constant is *τ*_*m,e*_ = 20 ms, and *R*_*m,read*_ = *τ*_*m,e*_/*C*_*m*_. Consistent with the idea that synapses projecting to other brain areas will also undergo depression, the dynamic weights *J*_*read,i*_(*t*) obey the same equation as all excitatory weights within the BCN.

To compute false positive and correct detection rates, a single lower decision boundary *θ*_−_ is used by the IR, as depicted in fig. 3. A detection event is registered if *A*_ir_ falls at least once below the boundary *θ*_−_ within the detection time window (0, *T*_*w*_ = 600 ms). In catch trials, no stimulus is present. These catch trials are used to determine the *false positive rate*:

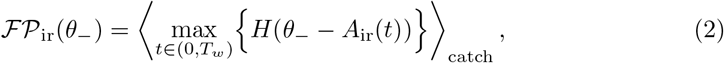

where *H*(*x*) is the Heaviside step function, and angular brackets indicate trial average. The reasons why a single lower detection boundary is used are explained below (see Firing-rate response of the network). The *hit* or *correct detection* rate is computed exactly in the same way, but in the presence of a stimulus

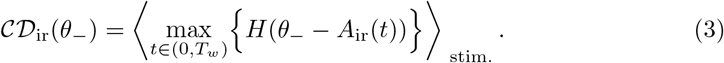

**Fig 3.**
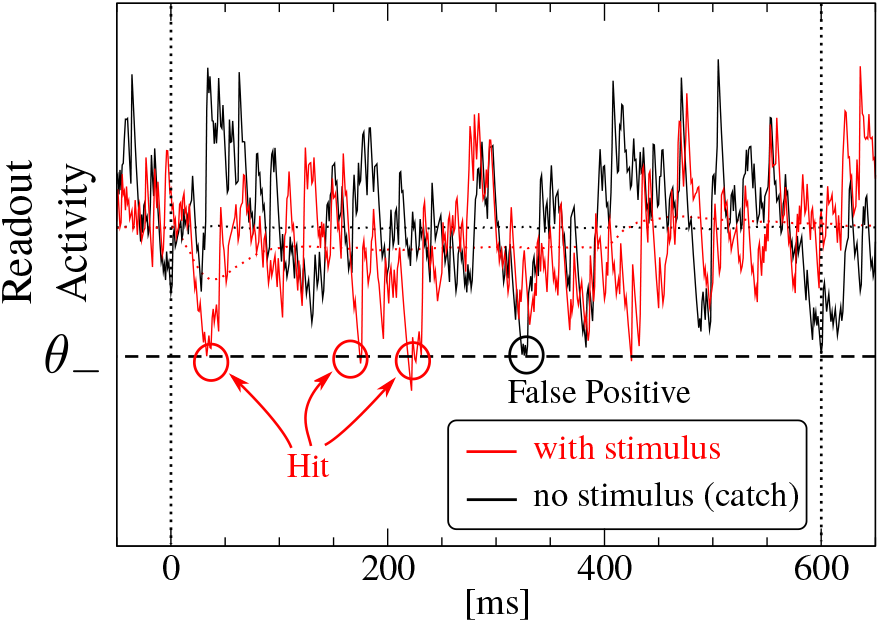
Working principle of the detector. Here, the case of a lower boundary is shown. The black trace is one realization of *A*_ir_(*t*) for a catch trial (no stimulus). Because the trajectory hits the detection boundary once, this trial generates a false positive event. The red trace represents one realization of the IR activity *A*_ir_(*t*) in the presence of a stimulus. Because the trajectory hits the boundary at least once (in this case even multiple times), this trial triggers a hit, or correct detection event.

#### Differentiator readout

The differentiator readout (DR) first reads in the input from the network in the same way as the IR and it takes the difference between *A*_ir_ evaluated at two times separated by a lag Δ*T*. The result is then convolved with an exponential filter 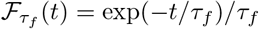 to reduce the noise

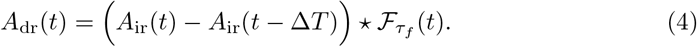

Trajectories computed from eq. (4) are used in combination with an upper detection threshold *θ*^+^ to obtain the false positive and hit rates:

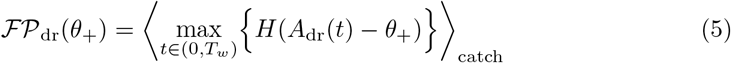

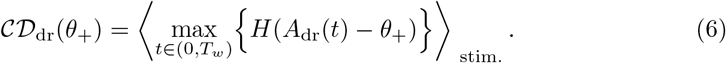

#### Differentiator network readout

The operation performed by the DR, i.e. the subtraction of a delayed copy of the readout activity, can be approximately implemented by the simple network shown in fig. 2C. The differentiator network readout (DNR) consists of two populations: one readout population of 10 000 RS neurons (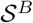) and one population of 2000 FS inhibitory neurons (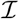). Both populations receive the same number of excitatory feed-forward connections from the RS population of the BCN. More precisely, each neuron in the DNR receives input from 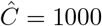 randomly chosen RS neurons. Neurons in the readout population 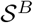 evolve according to the same dynamical equation as RS neurons of the BCN, while neurons in 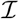 follow the same dynamical equations as the FS neurons of the BCN.

If the purpose of the DNR is to implement the operation performed by the DR, it is crucial that the feed-forward inhibitory pathway from the BCN via 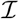 to 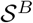 balances the direct feed-forward excitatory pathway input at a later time. To this end, the average weight of connections from 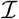 to 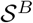, 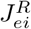 is chosen such that a static change in the input from the direct pathway Δ*μ*_*e*_ is compensated by the static change in the input from the inhibitory pathway 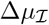 (see fig. 4). The value of 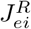 that approximately satisfies the condition 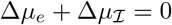 is computed from a linear-response calculation, reported in the Methods. Importantly, the inhibitory pathway is given an additional transmission delay Δ*T* = 10 ms, so that changes in the input from the BCN to the DNR are counterbalanced at a later time. More details on the DNR can be found in the Methods along with all parameter values.

**Fig 4.**
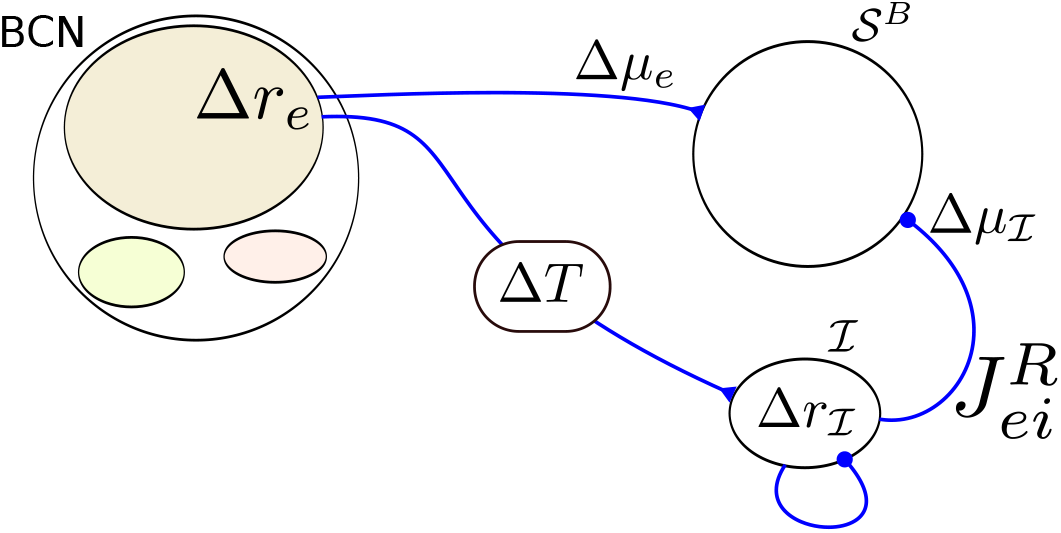
Tuning of the differentiator readout network to implement the operation of the differentiator readout scheme. A perturbation in the firing rate of the RS neurons in the BCN (Δ*r*_*e*_) causes a perturbation in the mean input to the RS readout neurons, Δ*μ*_*e*_, and a perturbation in the firing rate of the inhibitory readout population 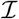. This change in firing rate causes a shift in the input from 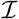 to 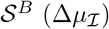. The strength of the connection from 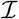 to 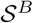 is adjusted such that 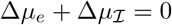. This cancellation reaches 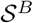 with a time lag Δ*T* so that the readout network roughly implements the operation of the DR eq. (4).

The DNR activity is obtained by filtering the average firing rate of the readout neurons 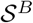 with the same exponential filter used for the DR:

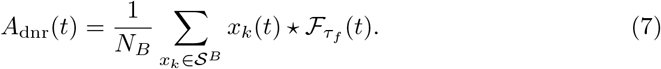

False positive and hit rates are obtained in exactly the same way as done for the DR:

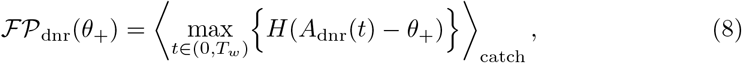

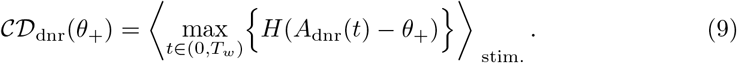

### Effect size

The *effect size* is defined as the difference between the hit and the false positive rate [4]. It is a function of the detection threshold *θ*

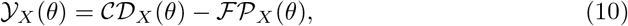

where *X* ∈ {ir, dr, dnr} indicates the detector type and *θ* can be either an upper or lower boundary. The false positive rate of 0.25 corresponds approximately to the average false positive rate measured experimentally. For this reason, this value was chosen to compare the simulation results to the experimental data. More precisely, the threshold 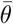 is chosen such that

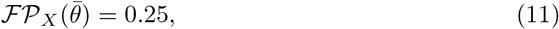

which is then used to compute

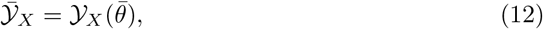

which is the final output of the detection procedure and will be compared to the experimental data.

### Single-cell stimulation

In every trial, the network is initialized with random initial conditions and simulated for *T*_idle_ = 1200 ms, to let the system forget the initial state. The spontaneous firing pattern of the network is asynchronous and irregular (fig. 1). The mean spontaneous firing rate of RS, FS, and SOM-LTS neurons is *r*_sp,e_ ≈ 0.8 Hz, *r*_sp,i_ ≈ 10 Hz, and *r*_sp,s_ ≈ 3 Hz, respectively. These properties of the spontaneous firing activity are consistent with experimental observations [45, 46].

A neuron is randomly selected as site of the nanostimulation, which is switched on at *t* = 0 and modeled as an additional current injected into the cell. The maximum stimulation current used here is Δ*I*_max,*e*_ = 5 nA. This value is chosen to elicit a similar number of spikes as in the experiment and is in the range used experimentally [47]. After the stimulus is switched off, the network is further simulated until the time reaches *t* = *T*_*end*_ = 1200 ms.

Following [6, 47], we use step currents of different lengths and intensities to investigate how the response probability depend on the properties of the stimulus. Furthermore, random permutations of steps of different length and amplitude will be used to generate irregular spike trains. Two equally-sized sets of catch trials, i.e. trials in which no stimulus was present, were simulated to estimate the size of random fluctuations in the detection rates.

The shot noise mimicking external input and the initial conditions were drawn anew in every trial. The same realization of the network (randomized cellular parameters and the connectivity matrix including weights and delays) was used once for each stimulus type. The total number of trials per stimulus type was *N*_trials_ = 10000.

### Firing-rate response of the network

Before investigating to what extent the three readout procedures introduced above are capable to detect the single-cell stimulation, it is instructive to examine the trial-averaged firing rate response of the network to the stimulation of a single RS cell. In the following, the case of a constant step current with intensity at 25% of the maximum and a duration of 400 ms is considered.

When a single RS cell is stimulated (red triangle in fig. 5), its output spikes affect a relatively small set of RS neurons (blue shaded area in fig. 5) because RS-to-RS connections are sparse (15% connection probability). Furthermore, their average amplitude is smaller if compared to other connections and they are strongly depressing, so that the direct effect on the overall firing rate of the RS population is small.

**Fig 5.**
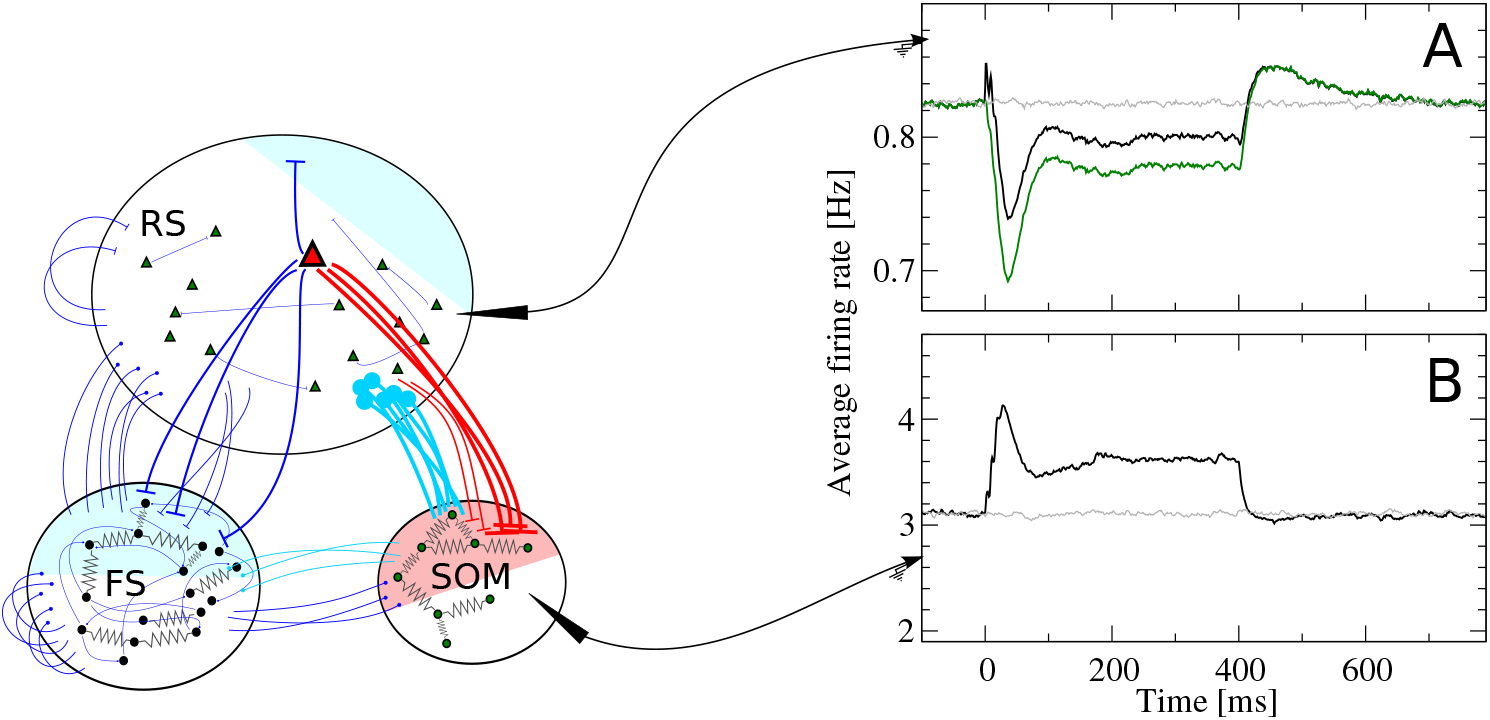
Disynaptic inhibition mediated by somatostatin-positive low-threshold spiking (SOM-LTS) cells causes inhibitory response to the stimulation of a regular spiking (RS) cell. **A**: when a RS cell is stimulated (the stimulus is switched here on at *t* = 0 and off at *t* = 400 ms), synapses from the stimulated cell (red triangle) to the SOM-LTS population strongly facilitate and cause a large increase in the firing rate of the SOM population, which then relaxes back to a plateau because of the spike-frequency adaptation (**b**). The inhibitory input from the SOM to the RS population produces a response in the RS cells which is almost a mirror image (**a**, black line). The initial positive peak in the RS response is due to the spikes fired by the stimulated cell itself, as it can be seen by excluding it from the population (**a**, green line). The curves in panels **a** and **b** are averages over 10000 trials and spikes are filtered with an exponential filter with decay constant *τ*_*f*_ = 15 ms. Gray lines indicate catch trials (no stimulation).

Connections from RS cells to the FS population are dense (40%), so that the spikes of the stimulated RS neuron reach a large fraction of the FS population (blue shaded area in fig. 5). However, FS cells also strongly inhibit one another and thus counteract the input from the stimulated cell. Consequently, the average change in the firing rate of the entire FS population is small (not shown).

Finally and most importantly, the output of the stimulated cell reaches a large fraction of the SOM-LTS population (50%, red shaded area in fig. 5) via strongly facilitating synapses. As a result, they induce an appreciable increase in the average firing rate of the SOM-LTS population, shown in fig. 5b. Importantly, SOM-LTS do not inhibit each other. However, the spike-frequency adaptation causes a strongly damped oscillation which, after an initial peak around *t* = 30 ms and a dip around *t* = 100 ms, relaxes to a plateau lying about 20% above the spontaneous firing rate level.

The increased inhibition from the SOM-LTS population ultimately causes the average firing rate of the RS population (fig. 5a, black line) to drop below the spontaneous level (fig. 5a, gray line). Note that the time course of the response of the RS population is closely related to that of the SOM-LTS population, except for the small peak shortly after the stimulus onset and for the overshoot after the stimulus is switched off. The small peak is due to the spikes fired by the stimulated cell itself. This can be seen by omitting the spikes fired by the stimulated cell (fig. 5a, green line) and observing that the peak disappears. The overshoot after *t* = 400 ms is due to the slow relaxation of the adaptation variable to its baseline value.

These observations are consistent with *in vitro* experiments showing that the strong activation of a single pyramidal cell in the barrel cortex has mostly an inhibitory effect on nearby pyramidal cells, and that this effect is due to disynaptic inhibition mediated by SOM-LTS neurons [40, 41]. More recent *in vivo* experiments pairing the stimulation of a single cell with calcium imaging suggest that bursts induced in a pyramidal cell have a very weak excitatory effect on other pyramidal cells and on FS neurons, but have a significant effect on neighboring SOM-LTS cells [48], in line with the behavior of our model.

### Relation between effect size and statistics of readout activity

In the previous subsection, the effects of the single-cell stimulation on the *trial-averaged* response of the RS population have been examined. The readout, however, must decide on the presence of the stimulus based on the RS population activity *in each single trial*, a much more difficult task (compare the smooth lines of fig. 5A,B with the noisy curves in fig. 3). The readout performance is quantified by the effect size, defined above. Before investigating how the effect size depends on the properties of the stimulus, we will examine how changes in the statistical properties of the readout activity *A*_*X*_ (*t*) can influence the effect size.

The simplest statistics that can be considered are the time-dependent mean and standard deviation of the readout activity (the averaging ensemble consists of the different trials). Statistics of higher order (skewness and kurtosis) were measured and did not display appreciable deviations from the spontaneous state and will be therefore omitted for brevity. Because we are interested in deviations from the spontaneous state, it makes sense to consider mean and standard deviation of the readout activity as standardized deviations from the spontaneous value. More precisely, we will consider first the time-dependent mean of the readout activity *A*_*X*_ (*t*) (here *X* =ir, dr, dnr, as defined in the subsection Readout models):

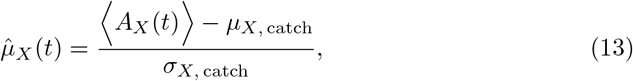

where *μ*_*X*, catch_ and *σ*_*X*, catch_ are the average mean and standard deviation in the spontaneous state, respectively:

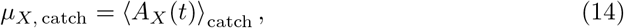

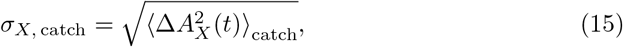

where 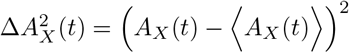, and the time dependence in both last equations is self-averaging due to the stationary conditions. The time-dependent standard deviation of the readout activity is defined in a similar way:

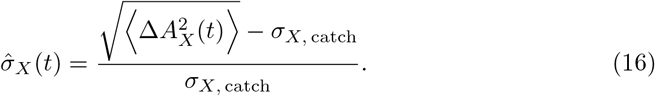

Non-zero values of 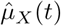 and of 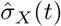 at any time point within the detection time window can impact the effect size in different ways. Suppose, for instance, that the considered detector employs an upper boundary. Then, a positive deflection of 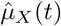 locally increases the probability of reaching the threshold, whereas a negative deflection reduces it. If a lower detection boundary is used, the opposite holds. Regardless of the type of threshold, a local increase of 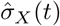 enhances the probability of reaching the threshold, whereas a local decrease in 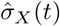 reduces the probability of crossing the decision barrier. This line of reasoning is qualitative only and holds under the assumption that *A*_*X*_ (*t*) is approximately normally distributed at all times.

To understand how multiple deviations from the spontaneous state within the decision time window jointly influence the effect size, it is useful to consider a simplified description of the decision model introduced in [25, 26, 49]. In this simplified theory, hit and false positive rates are approximated as the result of *n* = *T_w_/τ*_corr_ draws of a random variable, where *T_w_* is the detection time window and *τ*_corr_ is the autocorrelation time of the readout activity (in the example of fig. 6, *n* = 4). If these draws are treated as independent, the false positive rate reads

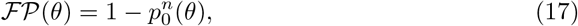

where *p*_0_(*θ*) is the probability of *not* crossing the barrier *θ* at a given time point and in the absence of the stimulus [*p*_0_(*θ*) does not depend on time and is therefore the same for each draw]. For concreteness, let us use an upper barrier at the value 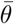, which yields the false positive rate of 0.25. In this way, the dependence on *θ* can be dropped, but the following considerations do not depend on the particular position or type of the boundary. Suppose now that 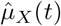 displays one peak at a certain position within the detection time window, as depicted in fig. 6A. Therefore, the probability that one trajectory of the readout activity triggers the detector is locally increased. Thus, in the vicinity of the peak, the probability of *not* triggering the detector will be *p*_1_ = *p*_0_ + Δ*p*_1_ < *p*_0_. The correct detection rate for this situation is then

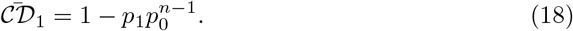

**Fig 6.**
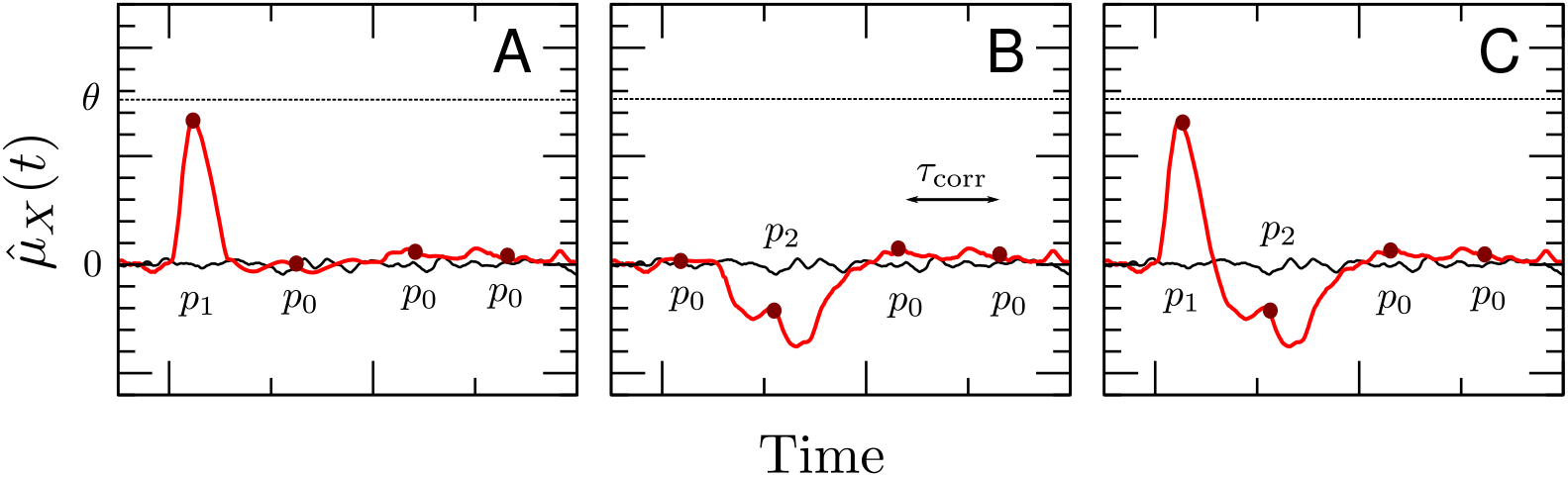
Illustration of simplified detection model used to interpret the simulation results. The continuous-time problem is approximated by a discretized process, obtained by “sampling” trajectories at times separated by the correlation time. Each deviation from the spontaneous state changes the probability of *not* reaching the decision barrier from the spontaneous value *p*_0_ to *p*_1_ or to *p*_2_.

Consequently, the effect size reads

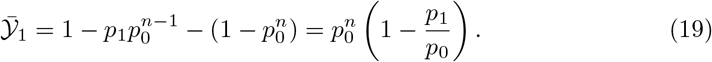

Consider now the situation of a negative deflection in 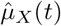 occurring at a different time (as in fig. 6B). Locally, the probability of not triggering the detector is *p*_2_ = *p*_0_ + Δ*p*_2_ > *p*_0_. In this case, the effect size is

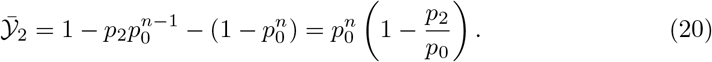

Suppose now that both features are present at sufficiently separated times within the same detection time window, as in fig. 6C. In this case, the effect size is

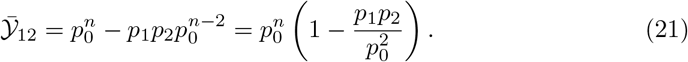

Substituting *p*_1_ = *p*_0_ + Δ*p*_1_ and *p*_2_ = *p*_0_ + Δ*p*_2_ into eq. (21) and supposing Δ*p*_1_, Δ*p*_2_ ≪ 1 yields

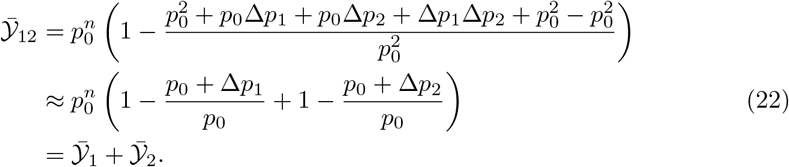

This approximation generalizes to the case of more than two deviations from the spontaneous state [49], given that all deviations are small compared to *p*_0_. For instance, when three features are present the effect size is

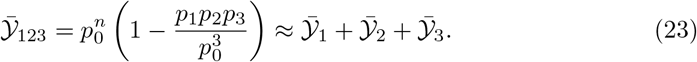

The main insight here is that weak deviations from the spontaneous state appearing in the same detection window can (approximately) add or cancel each other. This will be useful in the following to interpret how the behavior of 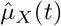 and 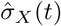 influence the response of the detector.

### Detection of the stimulation of a single neuron

In this subsection, we will investigate how the properties of the stimulus influence the response probability of the detector for each of the three readout procedures.

#### Effect of stimulus duration

The effect of changing the stimulus duration will be considered first. To this end, stimuli of length 100, 200, and 400 ms are used (fig. 7, top). The stimulus intensity is kept constant at 25% of the maximum current. In the experiment, when the stimulated cell was a RS neuron, the three stimuli evoked 6 ± 3, 11 ± 5, and 23 ± 10 spikes, respectively. In the model, the number of evoked spikes was 7 ± 1, 12 ± 2, and 20 ± 5 spikes, respectively. The average number of evoked spikes generated by the model is within one standard deviation of the experimental data. However, the spread of the spike count distribution is smaller in the model, which is not surprising, considering the multiple possible noise sources that are not modeled, and that only some of the cellular parameters are randomly distributed in the model.

**Fig 7.**
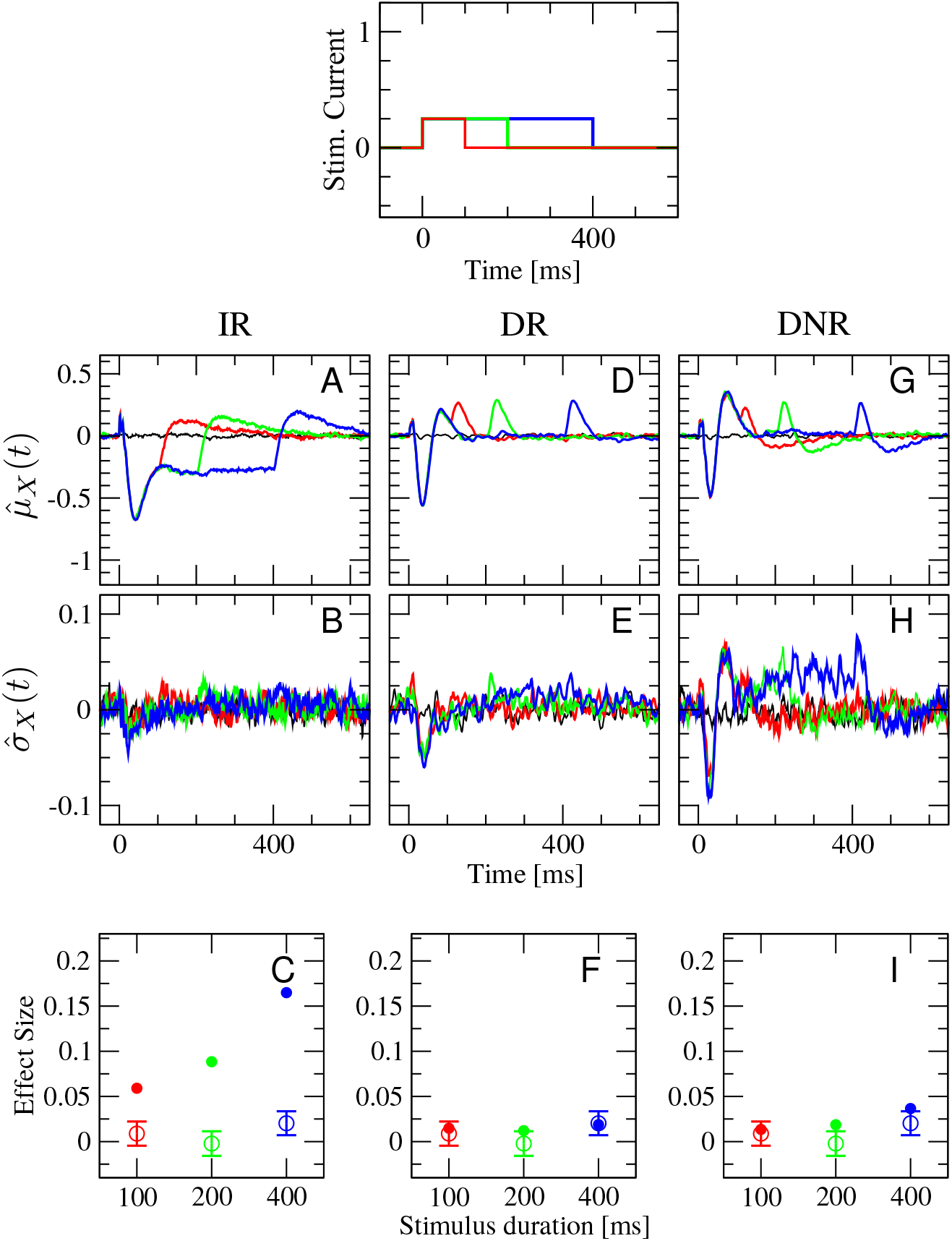
Only simulation results from differentiator readout (DR) and differentiator network readout (DNR) are compatible with the experimental data for the stimulation of a RS cell with three stimuli of equal intensity and different duration (top panel). First row: standardized deviation from the spontaneous value of the time-dependent mean readout activity eq. (13). Second row: standardized deviation from the spontaneous value of the time-dependent standard deviation of the readout activity eq. (16). Third row: average effect size for the three stimuli. Open circles with error bars are experimental results, which are the same in each panel, and represent the average effect size computed from 119 RS cells (2407 trials in total). First column: integrator readout (IR). Second column: differentiator readout (DR). Third column: differentiator network readout (DNR). The color of each line corresponds to a stimulus as in the top panel (red: 100 ms, green: 200 ms, blue: 400 ms). Black line is catch trial condition (no stimulus).

Figure 7 gives an overview of 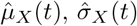, and the effect size measured by all three detectors with stimuli of different duration. In all plots, the color coding is as in the top panel and the black line represents catch trials. The various panels are organized as follows: the first, second, and third column represent results obtained from the IR, DR, and DNR, respectively; the first row shows the time-dependent trial average (expressed as standard score) 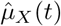, the second row shows 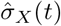, and the third row displays the effect size defined as in eq. (12) (filled dots) superimposed with the experimental results (open circles with error bars).

The IR activity *A*_ir_(*t*) is a low-pass filtered sum of a random subset of the spike trains fired by the RS population, as made clear by eq. (1). Hence, the time course of 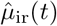 upon 400 ms stimulation (fig. 7A, blue line) closely resembles that of the RS population (see fig. 5 and the related discussion on p. 10). Except for the small initial peak due to the spikes fired by the stimulated cell (note that there is a 50% chance in each trial that the stimulated cell is included in the readout activity), the average response of the IR activity is a reduction that shows its deepest dip around *t* = 40 ms and then relaxes back owing to the spike-frequency adaptation of RS and SOM-LTS populations. In the case of the 100 ms stimulus (red line), the mean deviation goes back to zero shortly after the stimulus is turned off, while 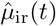 for the other stimulus durations (green and blue line) settles around −0.3 for the remainder of the respective stimulation time window. In each case, 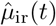 shows an overshoot caused by the slower relaxation time scales of the adaptation variables; the overshoot is stronger the longer the signal’s duration. The time-dependent standard deviation 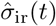, shown in fig. 7B, does not display appreciable deviations from zero except for a small dip that corresponds to that observed in the mean. The time courses of 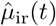 and 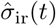 suggest that the readout activity has more chances of reaching a lower detection threshold for longer lasting stimuli. Indeed, the effect size measured by the IR strongly depends on the stimulus duration, as it can be seen in fig. 7C (filled circles), which is in contrast with the experimental data (open circles with error bars).

When the DR is used, the picture changes rather drastically. The time-dependent mean 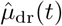 displays three peaks and one trough in response to all three signals (fig. 7D), even though the last two peaks partly overlap in the case of the 100 ms stimulus (red line). Because the DR considers differences in the IR readout activity, each peak corresponds to an upswing of 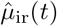 and the trough to the initial sharp drop. The most prominent feature in 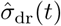 is the dip right after the stimulus onset, which is the same for all stimuli (fig. 7E). The main difference in the response to the three stimuli is the position of the last peak. Hence, it stands to reason that the DR activity *A*_dr_(*t*) has similar chances to reach the upper barrier regardless of the signal length. Indeed, the effect size measured by the DR is very similar for the three signals and of the same magnitude as that measured in the experiments (fig. 7F).

The time course of 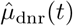 qualitatively resembles that of 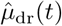 (see fig. 7G), thus suggesting that the DNR does approximately operate as a differentiator. The most evident difference between 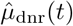 and 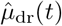 is the more pronounced undershoot after the last peak, which is likely due to the adaptation current in the readout population. The time-dependent standard deviation 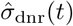 behaves similarly to the mean, and shows peaks and one dip at the same time as the time-dependent mean (fig. 7H). However, a persistent positive shift of 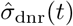 with respect to the zero level can be observed during the entire duration of the signal, which slightly enhances the detectability of the longer signal, as seen in fig. 7I. Still, the increase is moderate and the effect sizes are still close to the experimental values.

To summarize the results of fig. 7, the effect size measured by the IR is larger but strongly depends on the duration of the stimulus. The DR and the DNR detect the stimulus with a reliability that is essentially independent of the signal duration and that is of the same magnitude as the experimental data.

#### Effect of stimulus intensity

In the second experiment, we vary the firing rate of the stimulated cell by changing the current intensity while keeping the total area under the step stimulus, i.e. the injected charge, constant. As depicted in the top panel of fig. 8, the stimulus lasting 100 ms (red), 200 ms (green), and 400 ms have an intensity corresponding to 100%, 50%, and 25% of the maximum current, respectively. In this way, the total number of elicited spikes is approximately the same for each stimulus. In the experiment, the three stimuli evoked a firing rate of (109 ± 52) Hz, (54 ± 23) Hz, and (30 ± 10) Hz, respectively. In the model, the average evoked rates are (150 ± 25) Hz, (103 ± 20) Hz (50 ± 12) Hz. Note that the maximum current in the model is chosen such that the number of elicited spikes roughly matches the data of the previous experiment (dependence on stimulus duration), which were based on a *different* set of cells. Still, the evoked firing rates in the model are in a similar range as the experimental values.

**Fig 8.**
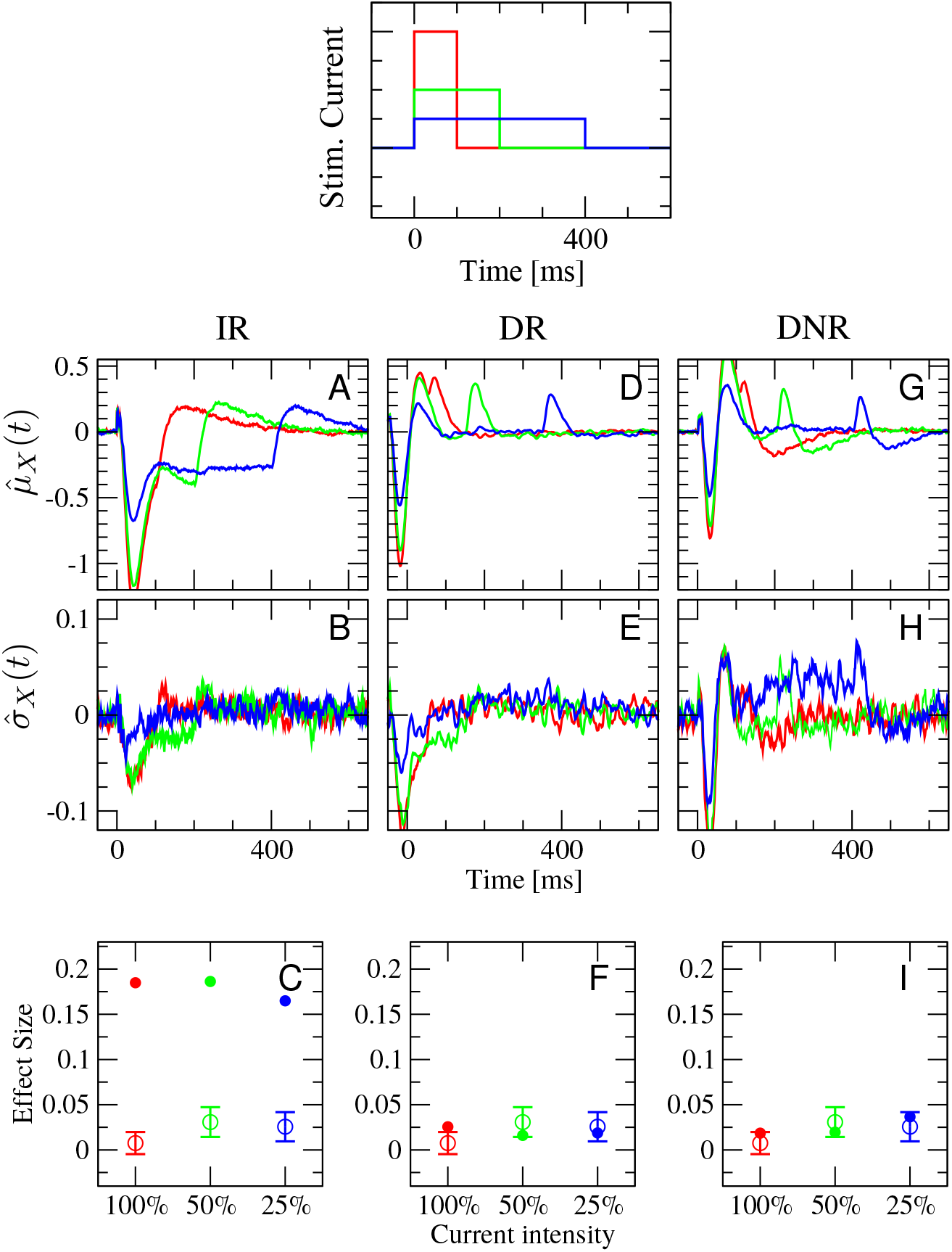
Only results from differentiator readout (DR) and differentiator network readout (DNR) are compatible with the experimental data for the stimulation of a RS cell with three stimuli of intensity inversely proportional to duration (top panel). As in the previous case, results from differentiator readout (DR) and differentiator network readout (DNR) are compatible with experimental data, whereas integrator readout (IR) gives qualitatively different results. Meaning of panels and color coding are as in fig. 7. Open circles with error bars are experimental results, which are the same in each panel, and represent the average effect size computed from 55 RS cells (1469 total trials).

Figure 8 reports all detection statistics arranged in the same way as before. The shape of 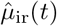, the mean response of the IR to the three stimuli, is similar to the previous case, although the initial drop is much stronger here when the stronger stimuli are used (fig. 8A). A clear dip in the time-dependent standard deviation 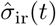 is observed right after the signal onset for the two stronger stimuli (fig. 8B). The pronounced initial response to the stronger signals roughly compensates the shorter duration of the signal in terms of chances of reaching the detection barrier. As a result, the effect size measured by the IR depends weakly on the current intensity and is much stronger than in the experiments, as shown in fig. 8C.

The time-dependent mean of the DR activity, 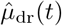, shown in fig. 8D, displays an initial drop followed by two peaks for each of the three stimuli. Here, however, both the first trough and the two subsequent peaks are more pronounced for signals of larger intensity, and the same holds for the initial dip in the time-dependent standard deviation 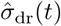 (fig. 8E). As these features have opposing effects and become stronger simultaneously upon growing stimulus intensity, the net effect on the response probability is barely noticeable, as shown in fig. 8F. Furthermore, the effect size is of magnitude comparable with the experimental observations.

Results obtained from the DNR are mostly similar to those from the DR. The principal differences are the mild trough after the last peak in the mean response 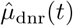 (fig. 8G) and the increased 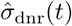 in the later part of the response (fig. 8H). The effect size measured from the DNR in the model shows a slight increase upon decreasing current intensity, which, considering the measurement uncertainties, is still consistent with the experimental observations.

Hence, the effect measured by all three readouts shows a weak dependence on the intensity of the stimulus, as it is observed in the experimental data. However, the effect size measured by the DR and the DNR is close to the experimentally observed average effect size of ≈ 2%, while the effect size obtained from the IR is considerably larger.

#### Effect of stimulus regularity

In the third and last *in silico* experiment, random stimuli will be used to evoke irregular spike trains. These stimuli, in accordance with the experimental procedure [6, 47], is a random shuffling of six current steps of length 10, 20, 40, 80, 160, and 90 ms, and with current intensity 100%, 50%, 25%, 12.5%, 6.25%, and −50% of the maximum current, respectively. In other words, each sequence consists of a random permutation of five positive (depolarizing) current steps with intensity inversely proportional to the duration and of one hyperpolarizing step, which inhibits the cell from firing. Two example signals are shown in fig. 9A. Note that stimuli are varied in each trial and not frozen. The irregular stimuli are constructed such that their total duration is 400 ms. The response probability to these stimuli will be compared to that of regular steps of 400 ms at 25% of the maximum current, which was used in both previous cases (plotted in blue). In the experiments, irregular current injections generated spike trains with an average firing rate of (24 ± 11) Hz and average CV of (1.1 ± 0.3). In the model, the average rate is (27 ± 5) Hz and the average CV (1.3 ± 0.3).

**Fig 9.**
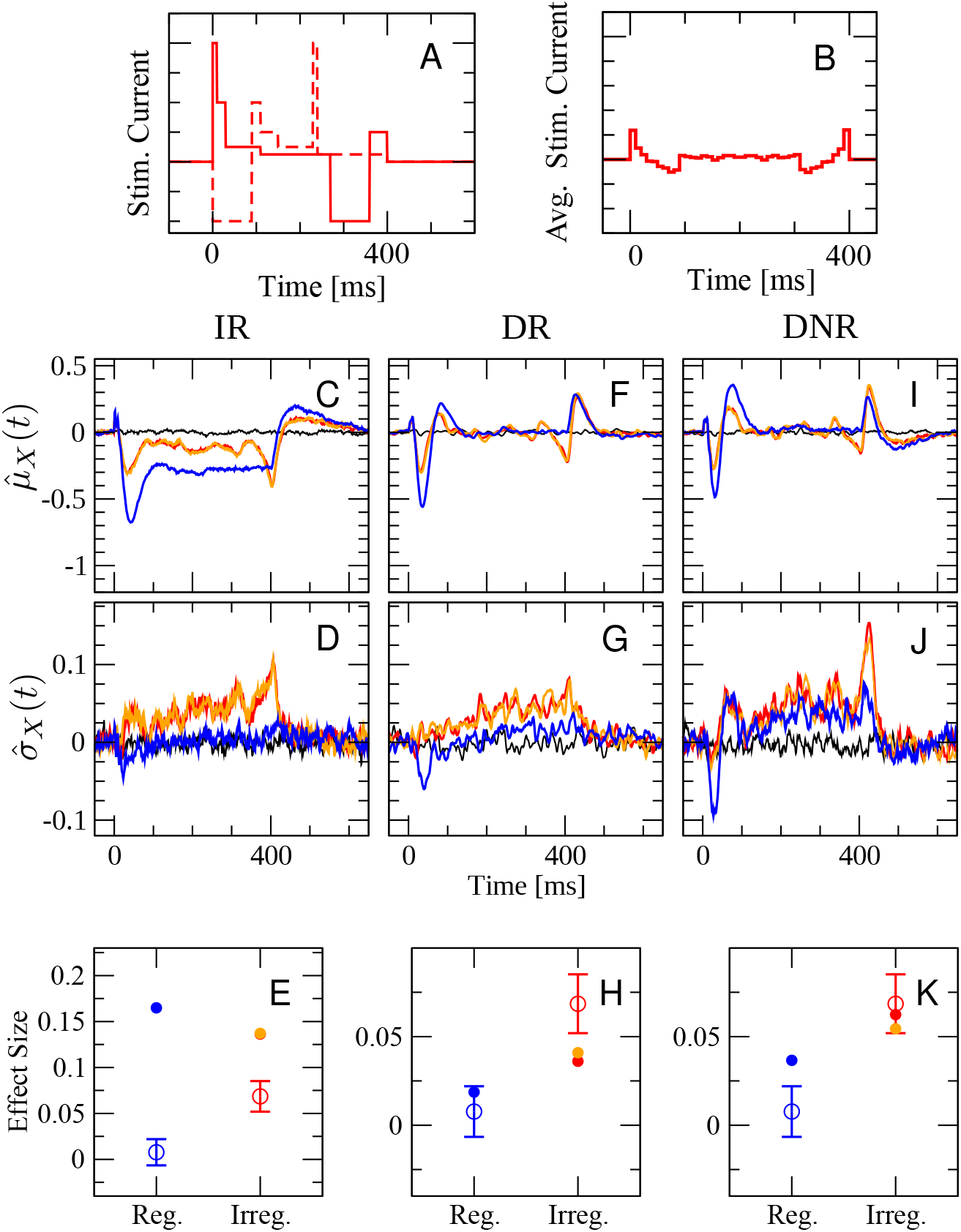
Summary of detection statistics upon stimulation of a RS cell with irregular stimuli (random permutation of multiple current steps), compared to regular 400 ms stimulation. Blue line refers to regular stimulus, as in the previous cases. Red and orange lines represent two different random samples of 10000 stimuli (the stimulus is changed in each trial) from the 720 possible permutations of the six steps (two specific realizations are shown in the top panel, while the average stimulus is shown in the inset). Black line is catch trial condition (no stimulus). First row: standardized deviation from the spontaneous value of the time-dependent mean readout activity eq. (13). Second row: standardized deviation from the spontaneous value of the time-dependent standard deviation of the readout activity eq. (16). Third row: mean effect size for regular and irregular stimulation. Open circles with error bars are experimental results, which are the same in each panel, and represent the average effect size computed from 62 RS cells (1780 trials in total).

In fig. 9 we compare the simulation results obtained when irregular stimuli are used to those obtained from the 400 ms regular current injection. The latter is shown once more in fig. 9 because it serves here as reference case but it will not be discussed in depth (see discussion above). Results for irregular stimulation are based on two sets of irregular stimuli, constructed by choosing from all possible permutations with equal probability. Despite the large total number of trials used here (*N*_trials_ = 10000), it is advisable to compare results for two independent sets of stimuli because the large number of possible permutations (6!=720) implies that the number of trials per signal is limited and that finite-size fluctuations due to the particular choice of signals may be non-negligible. Results for the two independent sets of irregular stimuli are plotted in red and orange in fig. 9. We recall that 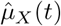 and 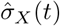 are obtained by averaging over different realizations of the irregular stimuli and are not related to the two particular signals shown in fig. 9A.

The *average* signal is plotted in fig. 9B and displays a peak just after the stimulus onset, followed by a mild trough and then by a plateau barely above the zero level. Just before the end of the stimulation time window, another trough followed by a peak can be seen. Accordingly, 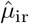 shows two dips at the same time where the two peaks in the average signal are seen (fig. 9C). Note that although the first and second half of the average signal are perfectly symmetrical, the second dip in 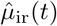 is more pronounced than the first.

Considering how different the average signal is from each particular realization of the irregular sequence, it may seem questionable to average over signals that provoke rather heterogeneous responses. However, the detector as well as the animals in the actual experiments do not know which realization of the irregular sequence is used in each trial. Therefore, this averaging ensemble, in which the stimulus is drawn in each trial, correctly represents the experimental situation and it makes sense to consider its time-dependent mean, as done above. The variability due to the particular realization of the input signal, which mostly averages out in the mean, reveals itself in an increased time-dependent standard deviation, 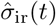 (fig. 9D), which is above the zero level in the entire stimulation time window and grows further towards the end of the stimulus. This increase of 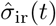 above the zero level enhances the chances of reaching the detection threshold. As a result, the effect size upon irregular stimulation is large, but not as large as that observed for the regular stimulus (fig. 9E filled dots), which is not consistent with the experimental observations, in which it is the other way around (fig. 9E, open circles with error bars).

The average DR activity in response to the irregular stimuli (fig. 9F, red and orange line) and to the 400 ms regular stimulus (blue line) are rather similar to each other. The main difference is that both the initial trough and peak are somewhat smaller for irregular stimulation. Furthermore, a small dip is observed for irregular stimuli just before the last peak. On the contrary, the standard deviation 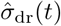 is markedly different for the two stimulus types (fig. 9G). When the regular stimulus is used, 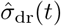 displays a small dip at the beginning and a slight increase in the later part of the stimulation time window (blue). In contrast, upon irregular stimulation it steadily grows over the entire stimulus range (red and orange lines), similarly to 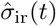. Consequently, the average effect size upon irregular stimulation measured by the DR is larger than that upon regular stimulation (fig. 9H, red and orange vs. blue full circles), which is qualitatively consistent with the experimental observations.

The time-dependent mean DNR activity 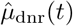 (shown in fig. 9I) is similar to 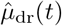, whereas 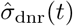 is generally larger than 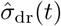, as in the previous cases. Nevertheless, fig. 9J shows that 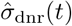 is larger in response to irregular stimuli (red and orange line) than its counterpart measured upon regular stimulation (blue line) and displays a strong peak at the end. The larger 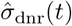 leads to a larger average effect size when irregular stimuli are presented (fig. 9K).

In summary, the simulation results in fig. 9 show that once again the DR and the DNR are consistent with the experimental observation that irregular stimuli are easier to detect than a regular current step, as opposed to the IR, which yields a smaller effect upon irregular stimulation than upon regular stimulation.

## Discussion

The development of juxtacellular stimulation has brought remarkable experimental opportunities, ranging from reliably evoking prescribed spike trains [50–52] to probing the role of a single neuron in the perception of sensory inputs and motor responses [2–6]. Recently, first attempts were made to model the behavioral responses to the stimulation of a single cortical neuron in rodents [25, 26]. However, how the behavioral response probability is influenced by the properties of the injected current [6, 47] is still unaccounted for theoretically. The aim of the present study was to construct a model that can reproduce some of these findings, i.e. that the probability of the behavioral response is not substantially influenced by the duration or by the intensity of a constant stimulus, but it is strongly dependent on whether irregular or regular stimuli are used.

The spiking network model we constructed in this study incorporates several features of the barrel cortex, and its parameters were consistent with the experimental literature. Among the biological processes included in the model, short-term depression and spike frequency adaptation could be expected to oppose slow changes in the input. However, our results indicate that these mechanisms may not be sufficient to explain the data if an integrator readout (IR) is employed. If a differentiator readout (DR or DNR) is used instead, simulation results are in agreement with the data.

How plausible is the scenario of a DNR and the implied transmission delays? The finite time difference used by the differentiator was chosen to be Δ*T* = 10 ms because it roughly matches the timescales of the changes induced in the readout population by the current jumps and thus ensures a good signal-to-noise ratio. The intersomatic distance required to achieve such a time difference in the barrel cortex would be approximately 2 mm [53], which is a large but not unphysiological value [54]. Moreover, although in the model the additional delay was entirely assigned to the connections from the BCN to the inhibitory readout population 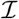 (fig. 2C), it would be possible to distribute the total latency among these connections and those from 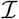 to the readout population 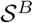. In this way, the disynaptic inhibitory pathway would need, for instance, to travel back and forth with a distance of only one millimeter.

It has been shown that a simple classifier can discriminate highly correlated inputs to a spiking network which exhibits chaotic spontaneous firing activity [55]. It is likely that some explicit training of the readout weights would drastically increase the effect size; in fact, the experimental subjects did undergo a training phase before the single-neuron stimulation sessions [4, 6]. However, we chose not to explicitly train the readout to detect specific cells, because the training in experiments was performed by employing microstimulation pulses, which intricately affect a large area rather than a specific cell directly [56–59]. Instead, here we assumed that the training had already occurred and that it resulted in the formation of the differentiator circuit. It is possible that training the readout to detect the microstimulation with a suitable learning rule would produce a detector that is also more efficient in detecting the single-cell stimulation than the simple differentiator considered here. It would be substantially more complicated, however, to construct a proper model for the complex and only partially understood cascade of events triggered by cortical microstimulation.

In our model, the main effect of inducing sustained firing in a single excitatory cell was to recruit SOM-LTS cells, which, in turn, inhibited the surrounding excitatory neurons. These results are consistent with the disynaptic inhibition observed *in vitro* [40, 41] and with recordings under anesthesia showing that bursts in pyramidal cells mostly activate surrounding SOM cells, hardly affecting other pyramidal cells or neighboring fast-spiking neurons [48].

Many different classes of inhibitory interneurons have been identified in the neocortex [60]. In this study we decided to include only two types of interneurons to try to limit the already high degree of complexity of the model. Modeling PV neurons is necessary as they are the most common interneuron type and form the backbone of the inhibitory system. SOM-LTS cells were included both because they are the second most common type of interneuron in the barrel cortex [61] and because other experimental studies hinted at their possible functional role when a single pyramidal cell is firing at high rates [40, 41, 48]. Another important class of cortical interneurons is formed by vasoactive intestinal peptide (VIP) neurons. These neurons do not directly provide inhibitory input to pyramidal cells and receive comparatively weak input from pyramidal cells. However, they have been found to make connections to and receive connections from SOM-LTS cells [32]. A recent computational study shows that the mutual inhibition between VIP and SOM-LTS cells can modulate the response of pyramidal cells to external input [62]. On this basis, it may be speculated that VIP neurons also amplify the response to the single-cell stimulation through disinhibition. In other words, SOM-LTS cell activation would inhibit VIP neurons which, in turn, would disinhibit SOM-LTS and amplify the effects of single-cell stimulation. Because VIP neurons are believed to receive top-down input, this conjecture would explain how the attention level of the experimental subjects positively influences the ability to detect the single-cell stimulation [4, 6].

The network model considered here represents the surroundings of the stimulated cells, which justifies the choice of a random unstructured connection topology within each neuronal population. Expanding the model beyond the local scale requires a structured or distance-dependent connectivity profile. Spatial connection profiles and non-random topologies have strong repercussions on the cross-correlations between the spike trains in a network [63–66]. Cross-correlations, in turn, largely contribute to the fluctuations in the pooled activity of a large readout population, as was considered here [25, 49, 67–70], and in general has consequences for the propagation of information about a stimulus to subsequent processing stages [71]. Hence, it is important that future studies investigate how different network topologies influence both the signal (the single-cell stimulation) and the noise (the fluctuations in the network’s activity).

Our results indicate that a readout circuit operating as a differentiator is in better agreement with the experimental data, although the effect size measured by the integrator readout was larger in all considered cases. Recalling that experimental subjects are rewarded for each correct detection, why should they opt for a sub-optimal readout procedure? One possible answer is that the integrator readout only works better for the stationary situation considered here. In reality, the detector would have to deal with strong and slow variations in the network’s firing state. It is possible to have a sense of this kind of variability by looking at fig. 10, which shows the spontaneous firing rate of some neurons during single-cell stimulation recording sessions. The strong changes in firing rate are mostly unrelated to any stimulation event. A readout that integrates spiking in a sliding time window would experience difficulty in distinguishing the possibly small changes induced by the single-cell stimulation from the strong background variations, whereas a differentiator readout might still separate the timescales of stimulus and background noise.

**Fig 10.**
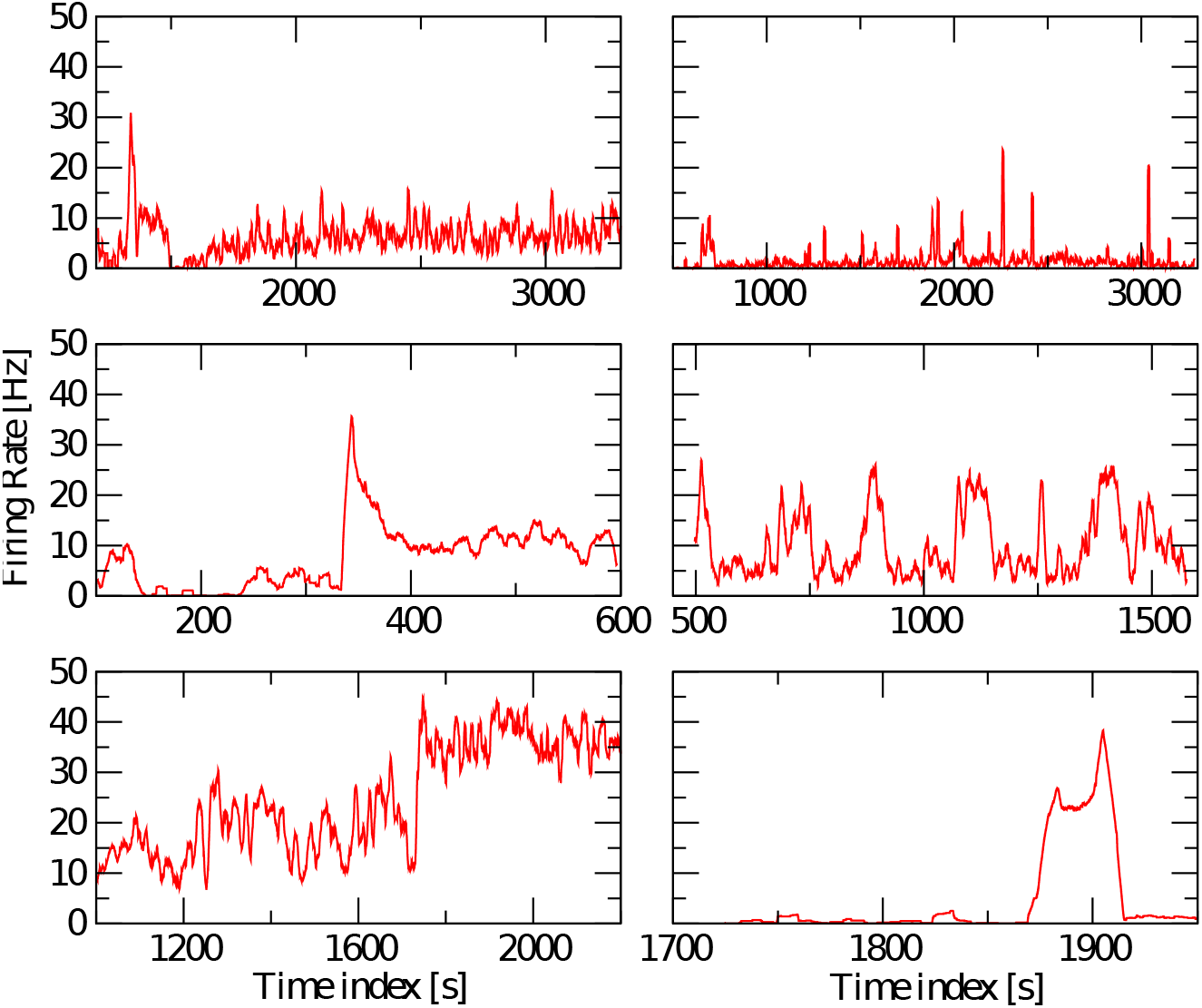
Nonstationarity of the firing rate of cells measured *in vivo* in the barrel cortex. Spikes are filtered with a 1 s sliding window.

## Methods

### Detailed description of the recurrent network model

#### Single-neuron properties and total input to neurons

We modeled all neurons as leaky integrate-and-fire point neurons [72]. The *k*th neuron follows the differential equation

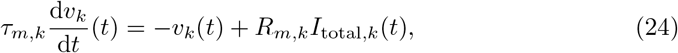

where the membrane time constant *τ*_*m,k*_ was drawn from a lognormal distribution with mean *τ*_*m,e*_ = *τ*_*m,s*_ = 20 ms if *k* is a RS neuron or a SOM-LTS neuron, or with mean *τ*_*m,i*_ = 10 ms if *k* is a FS neuron. The standard deviation of all three distributions was set to 20% of the mean. These values are compatible with experimental measurements for the rat barrel cortex [29, 30]. The membrane resistance is *R_m,k_* = *τ_m,k_/C_m_*, where the capacitance *C_m_* = 150 pF is assumed equal for all neurons. Equation (24) is complemented with the rule that whenever *v_k_*(*t*) reaches the threshold value *v_T,k_*, the neuron emits a spike and *v_k_*(*t*) is reset and clamped at *v_R_* = 10 mV for the duration of the refractory period *τ*_ref,k_. The value of the firing threshold was drawn independently for each neuron from a Gaussian distribution [30] with mean *v_T,E_* = *v_T,I_* = 20 mV if *k* is an RS or FS neuron [29, 30] and with mean *v_T,S_* = 14 mV if the *k*th neuron belongs to the SOM-LTS population, because the distance from resting potential to threshold is 5 mV to 7 mV lower in SOM-LTS neurons than in RS and FS neurons [29]. The standard deviation was set to 10% of the mean for all three neuron types [29]. The refractory time is 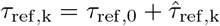, where *τ*_ref,0_ = 4 ms and 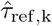 was drawn from a lognormal distribution with mean 2 ms and standard deviation 1 ms. The variability in the refractory time serves the purpose of mimicking the variability in the maximum firing rate of neurons [29].

If the *k*th neuron belongs to the FS population, its total input current *I*_total*,k*_ is just the sum of the external input and of the recurrent input, i.e. it reads:

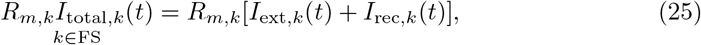

where the first term on the right side of eq. (25) represents the external input from other brain areas and the second term models the recurrent local input from other neurons within the network. If the considered *k*th neuron belongs either to the RS or to the SOM-LTS population, the total input current includes an additional adaptation term *a_k_*(*t*):

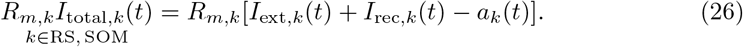

The adaptation current in the last equation obeys [36, 37]:

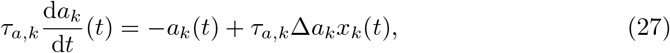

where *x*_*k*_(*t*) = ∑*_j_ δ*(*t* − *t_k,j_*) is the spike train emitted by neuron *k*. In other words, every time the neuron fires, the adaptation current jumps by Δ*a_k_*. Otherwise, it decays to zero with the time constant *τ_a,k_*.

Both Δ*a_k_* and *τ_a,k_* are randomly drawn from a lognormal distribution with standard deviation equal to 20% of the mean. For RS neurons, the mean of the two distributions are *τ_a,e_* = 100 ms and Δ*a_e_* = 0.3 nA, respectively; for SOM-LTS neurons they are *τ_a,s_* = 50 ms and Δ*a_s_* = 0.2 nA, respectively.

With this choice of parameters, the strength of the spike-frequency adaptation roughly agrees with *in vitro* measurements from the layer IV of the rat barrel cortex [29, 35].

#### External input to the network

The external input encompasses one constant term and two excitatory Poissonian shot-noise processes:

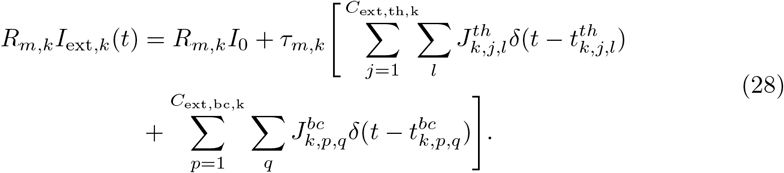

The constant term is set to *R_m,k_I*_0_ = 10 mV for all neurons. The second term in eq. (28) represents the input from the thalamus, and the third mimics incoming spikes from surrounding cortical areas. Because the thalamus has a higher average firing rate, “thalamic” input spikes at times 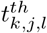 occur at an average rate of *r*_ext,th_ = 10 Hz, while “cortical” input spikes at times 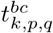 have a lower rate of *r*_ext,bc_ = 2 Hz. The number of external input spike trains depends on the cell type. Experimental studies suggest that SOM cells, in contrast to RS and FS cells, receive only weak input from the thalamus and from distant brain areas [29, 39]. Therefore, if the *k*th neuron belongs to the SOM-LTS population, then the number of external inputs is set to zero *C*_ext,th,k_ = *C*_ext,bc,k_ = 0, whereas when *k* is a RS or a FS neuron, then *C*_ext,th,k_ = 500. Furthermore, dendrites of FS neurons tend to be more localized than those of pyramidal cells, i.e. to receive more input from local RS neurons and less from distant ones. Hence, the number of inputs mimicking the cortical surroundings is *C*_ext,bc,e_ = 2000 when *k* is a RS neuron, and *C*_ext,bc,i_ = 1000 when *k* is a FS neuron. Each input spike causes a PSP drawn independently from an exponential distribution with mean *J*_ext,e_ = 0.1 mV when *k* is a RS neuron, and from an exponential distribution with mean *J*_ext,i_ = 0.2 mV when *k* is a FS neuron, because both thalamic and cortical excitatory postsynaptic potential (EPSP) amplitudes are larger in FS cells than in RS cells [29].

#### Recurrent input to RS neurons

The recurrent input term *I*_rec*,k*_(*t*) depends on the neuron type. If *k* is a RS cell, it is

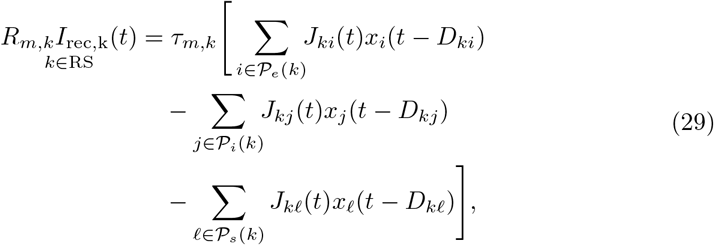

where *x_i_*(*t* − *D_ki_*) indicates the spike train emitted by neuron *i*, *D_ki_* represents the total transmission delay resulting from the axonal propagation, the neurotransmitter diffusion, and the dendritic propagation from neuron *i* to neuron *k*, *J_ki_* stands for the synaptic strength from neuron *i* to neuron *k*, which depends on the spiking history (see below).

Connections to neuron *k* originate from three sets of neurons: 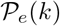, formed by *C_ee_* = 300 randomly selected RS neurons, 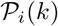, consisting of *C_ei_* = 200 randomly selected FS neurons, and 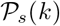, composed of *C_es_* = 100 randomly selected SOM-LTS neurons. Hence, the connection probability of RS-to-RS synapses is *C_ee_/N_e_* = 15%, of FS-to-RS and of SOM-LTS-to-RS is *C_ei_/N_i_* = *C_es_/N_s_* = 50%, consistent with the experimental observations that the connections between RS cells are sparse whereas those between RS and inhibitory cells are dense [27, 29, 40, 73–75]. Transmission delays are drawn uniformly in the range 0.5 ms to 1.0 ms [75]. All synaptic weights in eq. (29) undergo short-term depression (STD)

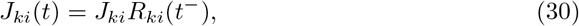

where the maximum coupling amplitudes *J_ki_* (corresponding to the first spike after neuron *i* has not been firing for a long time) are drawn independently from an exponential distribution with mean *J_ee_* = 0.1 mV for RS-to-RS connections, *J_ei_* = 0.5 mV for FS-to-RS coupling, and *J_es_* = 0.25 mV for SOM-to-RS connections. The variables *R_ki_*(*t*) represent the fraction of available synaptic resources, and *t*^−^ indicates that the function is evaluated immediately before a spike. Model and parameters of STD, i.e. the time evolution of the *R_ki_*(*t*), are described below.

#### Recurrent input to FS neurons

The recurrent input to a FS neuron reads:

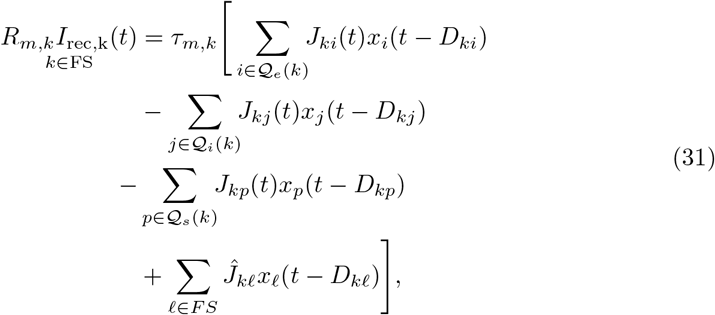

where the first term represents the synaptic input from 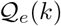, a set of *C_ie_* = 800 randomly selected RS cells (connection probability *C_ie_/N_e_* = 40%), the second term is the input from the inhibitory FS presynaptic population 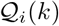 with size *C_ii_* = 200 (connection probability *C_ii_/N_i_* = 50%), and the third term represents the inputs from 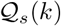, *C_is_* = 50 randomly selected SOM-LTS neurons (connection probability 25%). All weights appearing in these three terms follow eq. (30) and their peak value is drawn from an exponential distribution of mean *J_ie_* = 0.2 mV, *J_ii_* = 1.0 mV, and *J_is_* = 0.1 mV, respectively. Transmission delays are the same as for RS-to-RS connections. These values reflect the fact that FS neurons receive strong and dense connections both from RS and from FS neurons, and that synapses from SOM to FS neurons are comparatively weaker [29, 32]. The fourth and last term in eq. (31) is an effective model for the electrical coupling among FS cells (gap junctions), see next subsection.

#### Effective model for gap junctions

Both FS and SOM neurons in the rat somatosensory cortex are coupled by gap junctions [31, 33, 39, 76]. In a simplified picture, gap junctions act as a passive conductance coupling between the membrane voltage of two neurons. The standard way of mimicking the effect of a gap junction between neuron *k* and neuron *ℓ* would be the following additional current for neuron *k* [77, 78]:

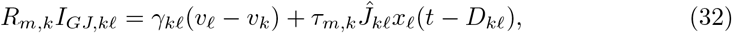

where *γ_kℓ_* is proportional to the Ohmic conductance between the two neurons and modulates the strength of the subthreshold coupling, and 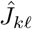 models the effect of spikes fired by neuron *ℓ*, which has to be added *ad hoc*, because LIF neurons do not explicitly generate action potentials. Gap junctions typically form between dendrites of different neurons. Therefore, the effect of spikes must travel from the soma along the dendrite of the first neuron to the gap junction and then from it along the dendrite into the soma of the second neuron. For this reason, the time necessary for this propagation can be as large as 0.5 ms [79]. Hence the delay term *D_kℓ_* is drawn from a uniform distribution in the range 0.1 ms to 0.5 ms. As reported in the main text, the subthreshold coupling is completely neglected here, i.e. *γ_jℓ_* = 0 is set for all neuron pairs. The subthreshold coupling was shown to have a very weak influence on the firing rate, synchrony, and oscillation frequency of networks of LIF neurons, as opposed to the spike-related coupling [34]. The amplitude of gap-junction-related post-synaptic potentials measured in FS neurons of the rat somatosensory cortex is rather variable and, on average, about half as large as excitatory post-synaptic potentials induced by RS neurons [79, 80]. Hence, 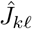 was drawn from an exponential distribution of mean 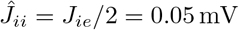.

The probability of a gap junction connecting two neighboring inhibitory neurons of the same type (FS with FS and SOM with SOM) is high (60% to 80% [39, 76]). For simplicity, the gap-junction coupling was approximated here as all-to-all (without self-coupling).

#### Recurrent input to SOM-LTS neurons

Finally, the recurrent input to a SOM-LTS neuron is

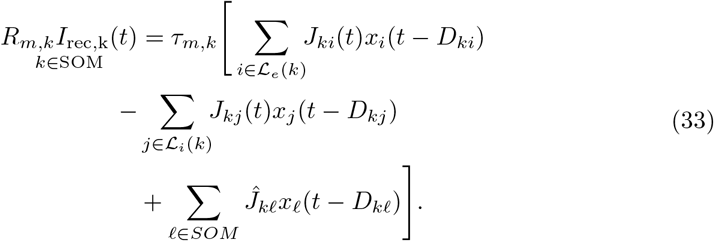

The three terms in eq. (33) represent the input from excitatory RS neurons, from inhibitory FS neurons, and from gap junctions, respectively. Gap-junctions are modeled in the same way as for FS neurons: their amplitudes and delays are drawn from the same distributions. The first term in eq. (33) is the input from *C_se_* = 1000 randomly chosen RS neurons (connection probability *C_se_/N_e_* = 50%). These are the only connections that undergo short-term *facilitation* instead of depression, and for which random *transmission failures* were modeled (details on the model below). The static baseline amplitudes *J_ki_* of each synapse are drawn independently from an exponential distribution and have mean *J_se_* = 0.1 mV. The second term in eq. (33) represents the input from *C_si_* = 100 randomly selected FS neurons (connection probability *C_si_/N_i_* = 25%). These connections have the average maximum strength *J_si_* = 0.25 mV, undergo short-term depression and obey eq. (30). Chemical synapses between SOM neurons are infrequent and weak [29, 39] and were omitted for simplicity.

#### Model of short-term depression

Except for those connecting RS to SOM neurons, all chemical synapses in the model undergo short-term depression. Each weight *J_kj_*(*t*) has a time dependence described by

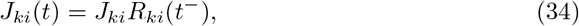

where the variable *R_ki_*(*t*) stands for the fraction of available synaptic resources and is described by the standard model by Tsodyks and Markram [43],

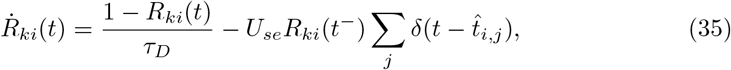

where 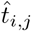 are the times at which the spikes of neuron *i* arrive at the synapse, and *t*^−^ indicates that the function is evaluated at *t* − *ε* (*ε* > 0 is a small positive number), i.e. just before a spike. The parameter *U_se_* represents the release probability and *τ_D_* is the recovery time scale. Note that the time evolution of *R_ki_*(*t*) depends on the spike times of the presynaptic neuron *i* only. Hence, if *τ_D_* and *U_se_* do not depend on *k*, the time course of each variable *R_ki_*(*t*) is a time-shifted copy of a single master variable *R_i_*(*t*)

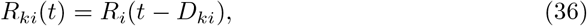

where *R_i_*(*t*) obeys the same equation as *R_ki_*(*t*), except that the arrival times 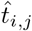 in eq. (35) are replaced by *t_i,j_*, the spike times of neuron *i*. Here, it is assumed that *τ_D_* and *U_se_* only depend on the type of the source and target neuron, but not on the identity of the particular neuron within a population so that eq. (36) holds. In this way, the actual number of dynamic variables required to simulate the network is reduced from one variable per synapse to one variable per neuron, which is an enormous computational advantage.

The parameter values chosen to model strong depression (all connections depicted in blue in fig. 1) are *τ_D_* = *τ_D,s_* = 150 ms and *U_se_* = *U_se,s_* = 0.2. With this choice, the eighth PSP of a 40 Hz pre-synaptic regular spike train is about one half of the maximal amplitude [29]. Most chemical synapses in the barrel cortex are depressing ([29, 42, 53]). However, inhibitory synapses originating from SOM-LTS neurons and terminating onto RS neurons show only weak depression or slight facilitation. Here, these connections are modeled as mildly depressing (fig. 1, light blue) by setting *τ_D_* = *τ_D,w_* = 50 ms and *U_se,w_* = 0.05. For simplicity, also SOM-to-FS connections were given the same STD parameters.

#### Short-term facilitation and transmission failures

Excitatory synapses from RS neurons to SOM-LTS neurons (depicted in red in fig. 1) are strongly facilitating ([29, 40, 41]). If the parameter *U_se_* in eq. (36) is turned into a dynamical variable, *u*(*t*), facilitating synapses can be described [44, 81]. The amplitude of the PSPs is proportional to the product *R*(*t*)*u*(*t*). Considering the connection from the RS neuron *i* to the SOM-LTS neuron *k*, the time evolution of the synaptic amplitude is described by (note that the conventions have been slightly changed with respect to ref. [44]):

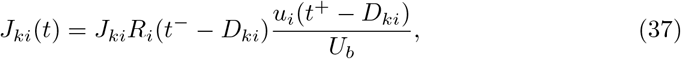

where *t*^+^ means that the function is evaluated at *t* + *ε*, i.e. the value of *u_i_*(*t*) just after the occurrence of a spike. The variables *R_i_*(*t*) and *u_i_*(*t*) obey

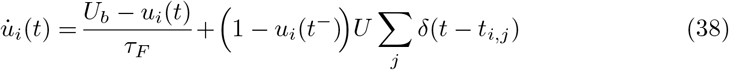

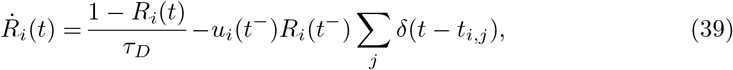

where *t_i,j_* indicate the spike times of neuron *i*. The first term in eq. (38) governs the relaxation of the facilitation variable to the baseline level *U_b_* and the second term determines a positive jump upon each pre-synaptic spike. The time evolution of the depression variable *R_i_*(*t*) has the same form of eq. (35), i.e. a purely depressing synapse, except that the release probability is the time-dependent function *u_i_*(*t*). The choice of the parameters *U, τ_F_, τ_D_* dictates whether, for a given firing rate, the synapse facilitates, depresses, or both [44]. Here, the four parameters appearing in eqs. (38) and (39) were set as follows: *τ_F_* = 300 ms, *τ_D_* = *τ_D,f_* = 100 ms, *U_b_* = 0.01, and *U* = 0.03. With this choice and for a pre-synaptic stimulation of 40 Hz, the synapse is purely facilitating [29].

RS-to-SOM synapses stand out from all other synapses considered here because of a much higher occurrence of transmission failures at low presynaptic firing rates (the average failure rate is ≈ 10% for RS-to-RS synapses, ≈ 5% for synapses to and from FS neurons, and ≳ 50% for RS to SOM-LTS synapses [29]). However, the failure rate of RS-to-SOM-LTS synapses decreases to ≈ 10% upon repeated stimulation at 40 Hz (failure rates for other synapses weakly depend on the presynaptic firing rate [29]). Here, transmission failures are modeled only for RS-to-SOM synapses via a stochastic binary variable *S*(*p_f_*):

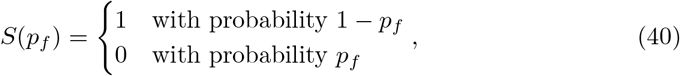

where *p_f_* describes the failure rate, which obeys the following dynamical equation

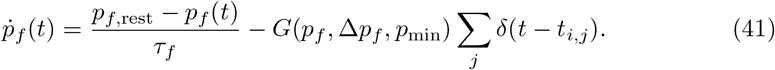

In the last equation, *p_f,_*_rest_ = 0.5 is the baseline failure rate. Upon each presynaptic spike, the failure rate decreases by *G*(*p_f_*, Δ*p_f_, p*_min_) and relaxes back to the baseline value with the time constant *τ_f_* = 250 ms. The size of each downward jump is Δ*p_f_* = 0.1 but is constrained to values above Δ*p_f_* = *p*_min_ = 0.1, a condition which is imposed by the piecewise linear function

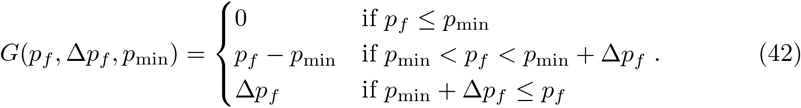

In the end, the synaptic weight from the RS neuron *i* to the SOM-LTS neuron *k* obeys the following equation:

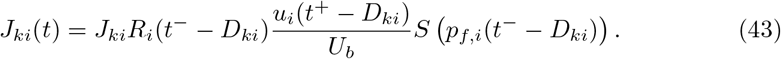

If the average effect of synaptic failures is taken into account, a 40 Hz presynaptic stimulation causes the eighth PSP to be about eight times larger than the first, which is in a reasonable qualitative agreement with the strong amplification measured *in vitro* [29, 41].

### Detailed description of the readout network model

The differentiator network readout (DNR) consists of one population of *N_B_* = 10 000 RS neurons 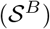 and of one population of 2000 FS neurons 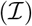. Each neuron in the DNR follows the same dynamical equation as its counterpart within the BCN and receives feedforward input from 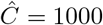 randomly selected RS neurons of the BCN. The number of independent Poisson processes mimicking thalamic input is 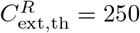 and the constant external input is *R_m_I*_0,_ = 15 mV. The number of independent Poisson processes representing cortical input is the same as for the BCN. Each neuron in the DNR receives 200 random connections from the local inhibitory population 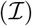. Hence, the only recurrent connections within the DNR are inhibitory.

All connections from the BCN to the DNR and within the DNR are randomly drawn from the same distributions as for the corresponding class of neurons within the BCN, except for the connections from the inhibitory readout population 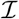 to the excitatory readout population 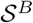, the average strength of which, 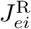, is tuned to a value that enables the DNR to approximate the function of a differentiator circuit, as explained in the following.

Referring to fig. 4, we have to calculate the value of 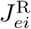 such that the input 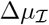 via the indirect path to the readout population 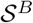 equals a negative and temporally delayed image of the direct input Δ*μ_e_*. This value can approximately be determined by the following linear-response calculation.

Consider a perturbation of the firing rate of the RS neurons within the BCN and indicate it with Δ*r_e_*. We assume that the perturbation is slow compared to the most important system time constants so that time-dependencies can be neglected. As a consequence of the firing rate perturbation within the BCN, the mean input from the BCN to 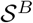 changes by

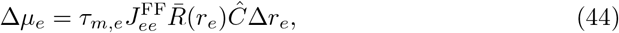

where the term

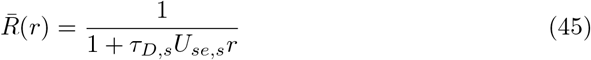

represents the average effect of the short-term depression (STD), given a presynaptic firing rate *r*. In eq. (44), 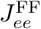 represents the average synaptic strength of the connections from the BCN to the excitatory readout population 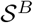. Likewise, the mean input from 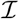 to 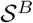 changes by

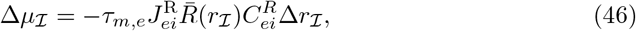

where 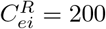 is the number of input connections from 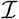 to 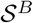 per postsynaptic neuron, and 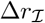 is the change in the firing rate of the population 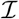 from the spontaneous value 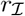.

The linear-response approximation of 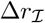 is

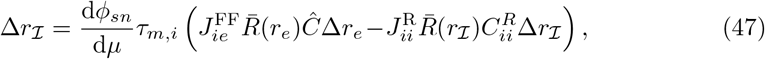

where d*ϕ_sn_*/d*μ* is the so-called DC susceptibility, i.e. the linear response of the firing rate of a LIF neuron to a slow change in its total mean input *μ*. The value of the DC susceptibility can be approximated by taking the derivative of the firing rate of a white-shot-noise-driven LIF neuron [82] with respect to its mean input. The explicit expression for d*ϕ_sn_*/d*μ* with a non-zero refractory period can be found in the first appendix of [49].

First, eq. (47) can be solved for 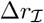 and substituted into eq. (46). Then, we require that the perturbation in the mean input from direct and indirect pathways cancel each other (see fig. 4). In other words, we impose 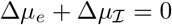 and finally solve for 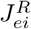, which yields

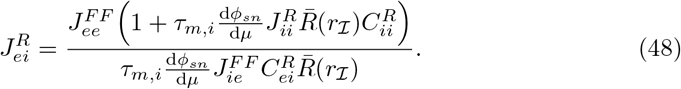

The only unknown quantity on the right hand side of eq. (48) is 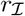, the spontaneous firing rate of 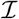. This firing rate can be estimated from the numerical solution of the following self-consistency condition:

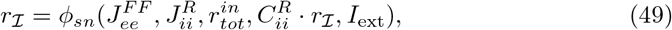

where 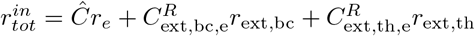 is the total excitatory input rate to 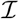 and *ϕ_sn_*(*a_e_, a_i_, R_e_, R_i_, I*_0_) is the firing rate of a LIF neuron driven by white shot-noise with exponentially distributed weights [82]. The first two arguments, *a_e_, a_i_* are the excitatory and inhibitory mean input weights, respectively. The third and fourth argument *R_e_, R_i_* are the input rates of the excitatory and inhibitory input, respectively. The last argument *I*_0_ is the constant input. The explicit expression with non-zero input current and non-zero refractory period is

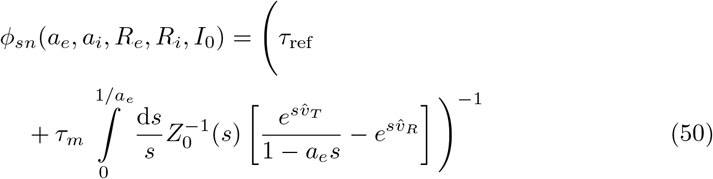

where 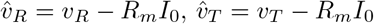, and 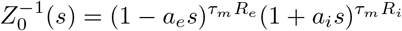.

Substituting numerical values in eq. (48) reveals that 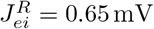 approximately satisfies the imposed condition.

### Experimental data

The experimental data appearing in figs. 7 to 9 are a part of the dataset of references [6, 47]. In particular, the data shown in fig. 7 are the average effect size (for each stimulus duration) of most cells shown in Fig. 18A of reference [47] (800 ms stimuli were not used in the present study); the experimental effect size in fig. 8 is the average for each stimulus intensity (and duration) of the cells appearing in Fig. 14A of reference [47]; the average effect size for regular and irregular stimulation of fig. 9 is based on the same dataset used for Fig. 21C of reference [47]. For experimental procedures, we refer to [6, 47].

## Acknowledgments

This work was supported by the German Research Foundation (DFG), GRK 1589/2 and by the German Federal Ministry of Education and Research (Bernstein Center II, grant no. 01GQ1001A).

